# The septate junction protein Snakeskin is critical for epithelial barrier function and tissue homeostasis in the Malpighian tubules of adult *Drosophila*

**DOI:** 10.1101/2020.12.14.422678

**Authors:** A.J. Dornan, K.A. Halberg, L.-K. Beuter, S.-A. Davies, J.A.T. Dow

**Author notes:** Correspondence to JATD. These authors contributed equally to this work (A.J. Dornan, K.A. Halberg).

## Abstract

Transporting epithelia provide a protective physical barrier while directing appropriate transport of ions, solutes and water. In invertebrates, epithelial integrity is dependent on formation, and maintenance, of ‘tight’ septate junctions (SJs). We demonstrated that *Drosophila* Malpighian (renal) tubules undergo an age-dependent decline in secretory transport capacity, which correlates with mislocalisation of SJ proteins and coincident progressive degeneration in cellular morphology and tissue homeostasis. By restrictively impairing, in adult tubules, the cell adhesion protein Snakeskin, which is essential for smooth SJ formation, we observed progressive changes in cellular and tissue morphology that phenocopied these effects, including mislocalisation of junctional proteins with concomitant loss of cell polarity and barrier function. Resulting in significant accelerated decline in tubule secretory capacity and organismal viability. Our investigations highlight the tubule’s essential role in maintenance of organismal health, while providing measurable markers of compromised epithelial barrier and tissue function that manifest in advanced morbidity and death.

Model for epithelial dysfunction arising from failure of smooth septate junctional complexes as a consequence of impaired *Snakeskin* expression.

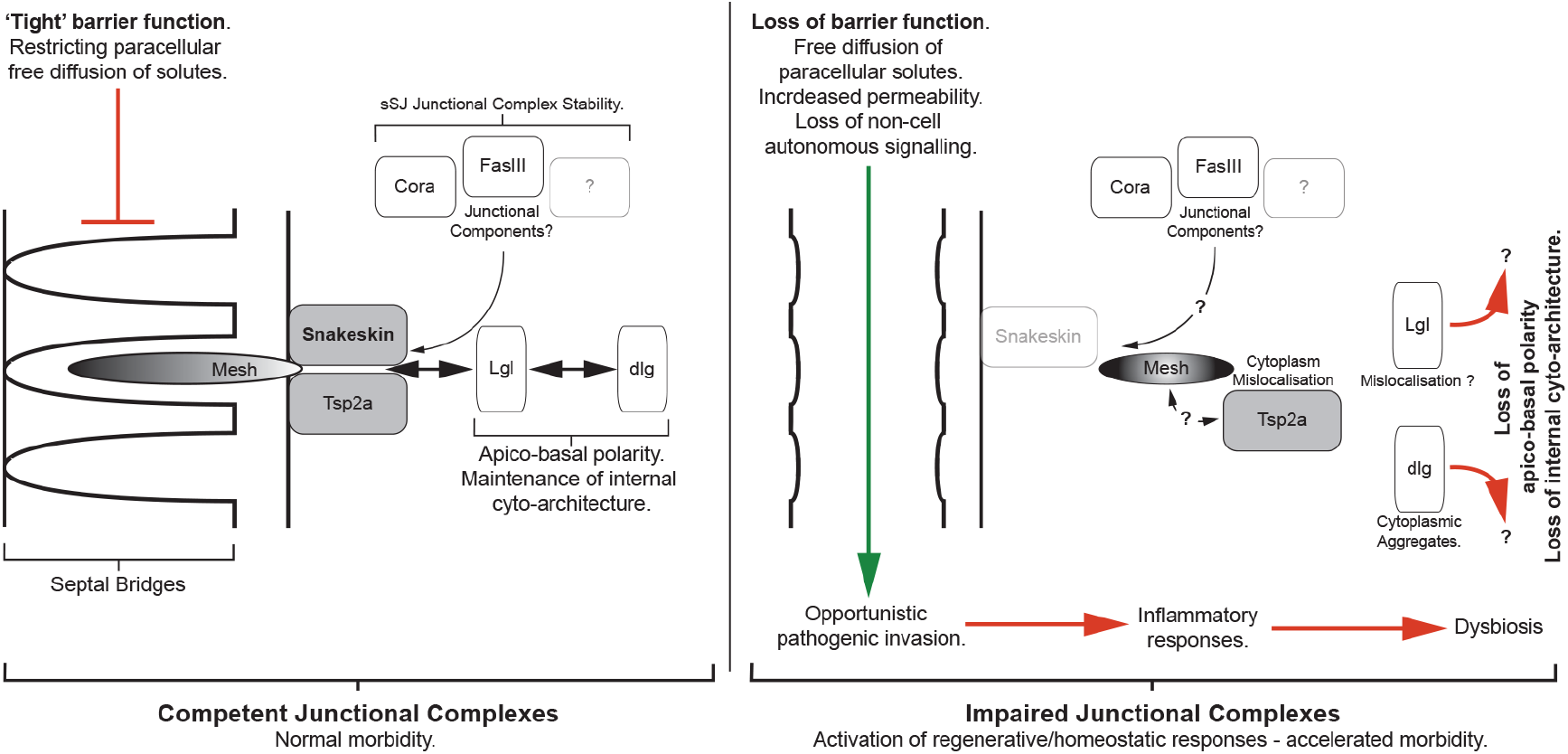

## INTRODUCTION

In multicellular animals, ageing presents as a progressive decline in tissue homeostasis and organ function, leading to increasing probability of disease and death (Rera et al., 2013b; Rose et al., 2012). Regulation of tissue homeostasis is thus critical to organismal lifespan; yet the molecular and cellular activities responsible for mediating age-dependent changes in tissue function, and how this in turn impacts organismal longevity, remain largely unexplored.

Maintenance of a healthy intestine has recently emerged as a critical determinant of lifespan across taxa, with stereotypic hallmarks of intestinal ageing including augmented stem cell behaviour, blocked terminal differentiation, activation of inflammatory pathways, reduced nutrient uptake, loss-of-barrier integrity and dysbiosis (Clark and Walker, 2018; Clark et al., 2015; Davies et al., 2012; Hu and Jasper, 2017; McGee et al., 2011; Regan et al., 2016; Rera et al., 2013a; 2012; Resnik-Docampo et al., 2017; 2018; Salazar et al., 2018). Notably, a causal link between these phenotypes and age-related remodelling of cell-to-cell junctions has been established in proliferative tissues such as the intestine (Clark et al., 2015; Izumi et al., 2019; Resnik-Docampo et al., 2017; 2018; Salazar et al., 2018), indicating that dysregulation of junctional proteins in self-renewing tissues may be a principal driver of ageing. However, whether age-related changes in cell-to-cell junctions also occur in tissues defined by low or no cell-turnover, like the nervous system, heart and kidneys, and whether such alterations significantly contribute to tissue degeneration and/or age-onset remains unresolved.

In insects, the renal (Malpighian) tubules (MTs) constitute the functional analogue of the vertebrate kidneys (Cohen et al., 2020; Dow et al., 2018) and are, as in vertebrates, considered to be a non-proliferative tissue (Skaer, 1993). The MTs are the principle organs responsible for maintaining water and ion homeostasis, yet serve additional roles in xenobiotic detoxification (Yang et al., 2007) and immunity (Davies et al., 2012; Verma and Tapadia, 2014). In the fruit fly *Drosophila melanogaster*, the MTs are comprised mainly of two physiologically distinct secretory cell-types; the principal (PC) cell and intercalated or ‘stellate’ (SC) cell (Beyenbach et al., 2010; Dow, 2012; Sözen et al., 1997). PCs are the sites of active cation transport, energised by V-ATPases localized apically to a prominent brush border (Halberg et al., 2016), whereas the smaller SCs control channel-mediated Cl^−^ and water fluxes (Beyenbach et al., 2010; Cabrero et al., 2020; Dow, 2012). Primary urine production in *Drosophila* is achieved through the integrated actions of PCs and SCs, driving transepithelial transport of ions and water. A third population of ‘tiny’ cells, found only in the ureter and lower tubule, have been proposed to be renal stem cells (Martínez-Corrales et al., 2019; Singh and Hou, 2009; 2008; Singh et al., 2011; Takashima et al., 2013; Wang and Spradling, 2020), yet to what extent these potentially proliferative cells contribute to tissue repair/homeostasis is unknown.

In invertebrates, epithelial integrity is maintained by either pleated or smooth septate junctions (pSJs and sSJs respectively), lateral intercellular contacts that are functional analogues of vertebrate tight junctions (Furuse and Izumi, 2017), with sSJs the dominant junctional complex in the MTs (Beyenbach et al., 2020; Jonusaite et al., 2020; Lane and Skaer 2016; Noirot-Timothee and Noirot, 1980). The putative cell adhesion molecule Snakeskin (Ssk), along with its counterpart Tetraspanin 2a (Tsp2a) and the membrane spanning protein Mesh are critical for the proper formation of sSJ’s (Beyenbach et al., 2020; Izumi et al., 2019; 2016; 2012; Jonusaite et al, 2020; Salazar et al., 2018; Xu et al., 2019; Yanagihashi et al., 2012). Mesh/Ssk/Tsp2a are also necessary, both individually and as a complex, for the appropriate localisation of associated junctional proteins such as Lethal giant larvae (Lgl) and Discs large (Dlg), both part of the Scribble polarity module (Khoury and Bilder 2020), Coracle (Cora) and Fascilin III (FasIII) (Chen et al., 2020; Izumi et al., 2012; 2016; Salazar et al., 2018; Yanagashi et al., 2012).

Here, we demonstrate for the first time that the adult MT undergo an age-dependent decline in secretory transport capacity, which correlates with mislocalisation of septate junction proteins and coincident progressive degeneration in cellular morphology and overall tissue homeostasis. We further show these effects are phenocopied by acute loss of *Ssk* expression, in either PC or SC sub-populations, of adult MTs. *Ssk* impairment in either cell population resulting in mislocalisation of junctional components, again manifesting in overt degeneration in cellular and tissue morphology. Critically, this acute failure of junctional integrity leads to an accelerated reduction in secretory capacity and concomitant loss of systemic fluid homeostasis, which ultimately results in a significant reduction in organismal lifespan. Furthermore, cell-specific manipulations of *Ssk* expression led to a pronounced increase in SC clustering, a block in SC maturation and a loss in apicobasal polarity, indicating a key role for *Ssk* in maintaining MT function and stability. Finally, knocking down *Ssk* expression also led to a striking proliferation of tiny (renal stem) cell as well as a dramatic increase in tubule tracheation, suggesting that MTs can autonomously respond to tissue damage and that Ssk acts as a novel regulator of tissue homeostasis in the tubule.

Taken together, our work demonstrates a crucial link between cell-cell junction integrity, epithelial transport competence and tubule homeostasis in a classically non-proliferative tissue, which provide novel insights into the mechanisms underlying tissue degeneration and age-onset.

## RESULTS

### Cell-specific Ssk depletion impairs systemic osmoregulation and reduced organismal lifespan

Failure to form sSj’s properly in early development, either ubiquitously or restricted to the MTs, is lethal (Beyenbach et al., 2020; Jonusaite et al., 2020; Izumi et al., 2019; Yanagihashi et al., 2012). We therefore employed temperature-sensitive tubulin GAL80 (McGuire et al., 2004), in conjunction with either a SC-specific, *c724*GAL4 (Sözen et al., 1997), or PC-specific, *Uro*GAL4 Terhzaz et al., 2010), GAL4 driver acting on UAS-*Ssk*^RNAi^ (Yanagihashi et al., 2012) to restrictively knock down *Ssk* expression in adult tubules (Figure S1A) (full genotypes see Methods). At the permissive temperature (18°C), GAL4 expression is repressed and *c724*^*ts*^>*Ssk*^RNAi^ and *Uro*^*ts*^>*Ssk*^RNAi^ (respectively designated as SC^*Ssk*RNAi^ and PC^*Ssk*RNAi^) flies develop into viable, fertile adults (Figures 1A and S1A). However, when transferred to the restrictive temperature (29°C) at the pre-pupal stage GAL4-drives expression of *Ssk*^RNAi^ in a tissue and developmentally restricted manner (Figure S1A-B). These SC^*Ssk*RNAi^ and PC^*Ssk*RNAi^ experimental animals successfully eclosed as adults. However, over time they progressively developed a ‘bloated’ phenotype due to increased water content (Figure 1A), a clear indicator of compromised MT epithelial function which resulted in significantly reduced viability (Figure 1B).

**Figure 1.**
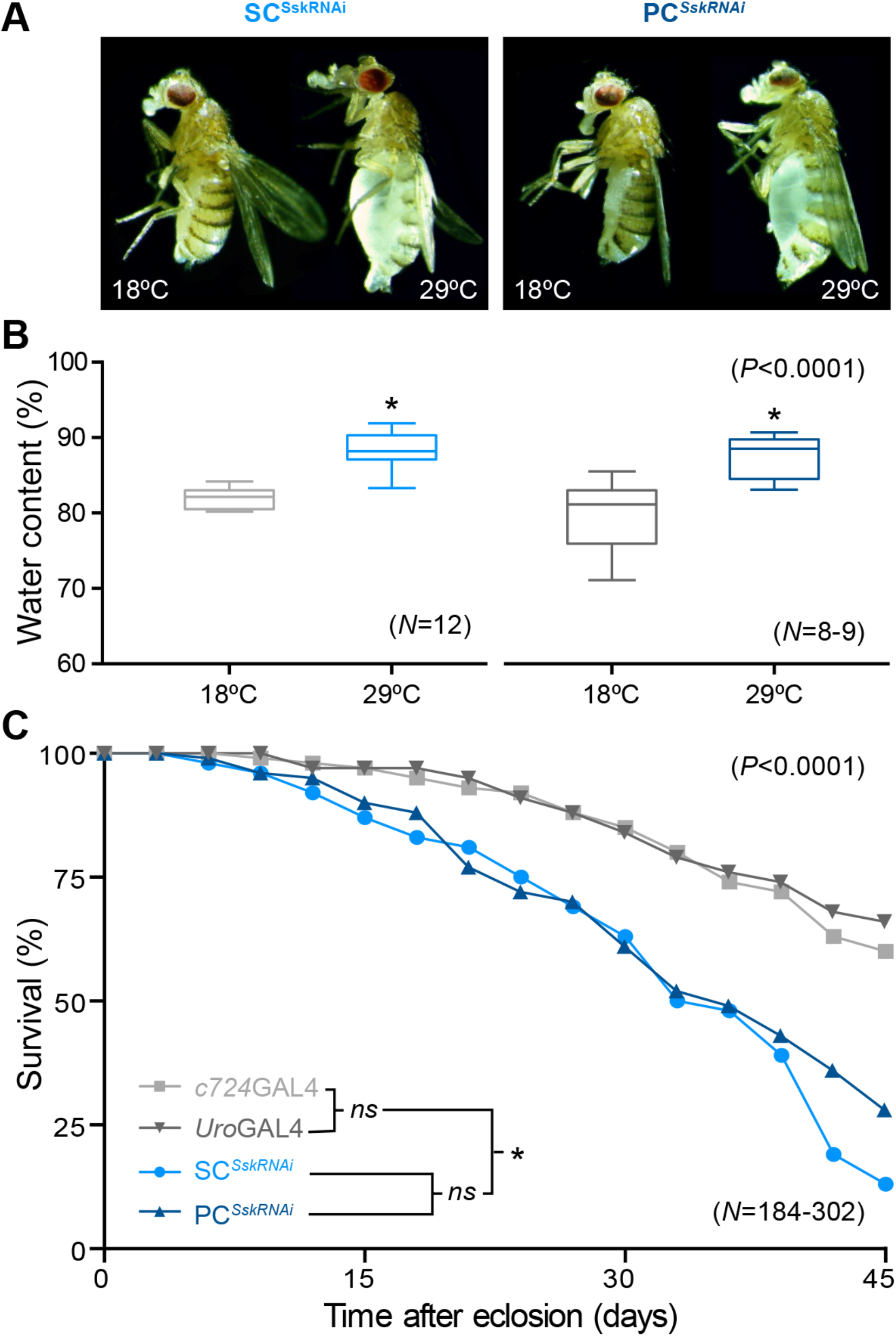
Cell-specific *Ssk* depletion in adult Malpighian tubules results in loss of fluid integrity and significantly reduced viability. (A) SC^*Ssk*RNAi^ and PC^*Ssk*RNAi^ adult females raised at the restrictive temperature (29°C) exhibiting an overt bloated abdomen phenotype as compared to controls raised at the permissive temperature (18°C). This ‘bloated’ phenotype is reiterated in males. (B) Comparison of ‘wet’ to ‘dry’ weight differences for experimental (29°C) female SC^*Ssk*RNAi^ and PC^*Ssk*RNAi^ 5 D adult females versus (18°C) controls. (C) Survival assay demonstrating significant reduction in viability of experimental SC^*Ssk*RNAi^ and PC^*Ssk*RNAi^ adult flies at 29°C versus controls. Student’s T-Test. P<0.0001. N, parentheses; wet-dry assay, each individual set=20 flies.

### Cell-specific Ssk depletion compromises cellular morphology

We then set out to determine what specific cellular deficits occur when Ssk is depleted in adult MTs that might contribute to this compromised epithelial function. When *Ssk* is knocked down in either SCs or PCs of adult tubules, junctional complex organisation – as realised by staining for Discs large (Dlg), a junctional protein required for structure, cell polarity, and proliferation control in epithelia (Bilder et al., 2000; Khoury and Bilder, 2020; Woods et al., 1996) – is overtly compromised compared to the more regular distribution exhibited in controls (Figure 2A-B), with mislocalisation of Dlg apparent as accretions in the cytoplasm (Figures 2B-C and S2B-C). Notably, SCs from SC^*Ssk*RNAi^ animals failed to develop their stereotypic mature ‘stellar’ morphology, instead appearing cuboidal and extruded from the tubule (Figure 2B-C). This extrusion likely results from development of an internal vacuole, which dramatically increases cellular volume (Figure 2D). Over time SC^*Ssk*RNAi^ and PC^*Ssk*RNAi^ mutant cells undergo apoptosis, losing GFP and Dlg expression and nucleation (Figures 2B-C and S2B-C).

**Figure 2.**
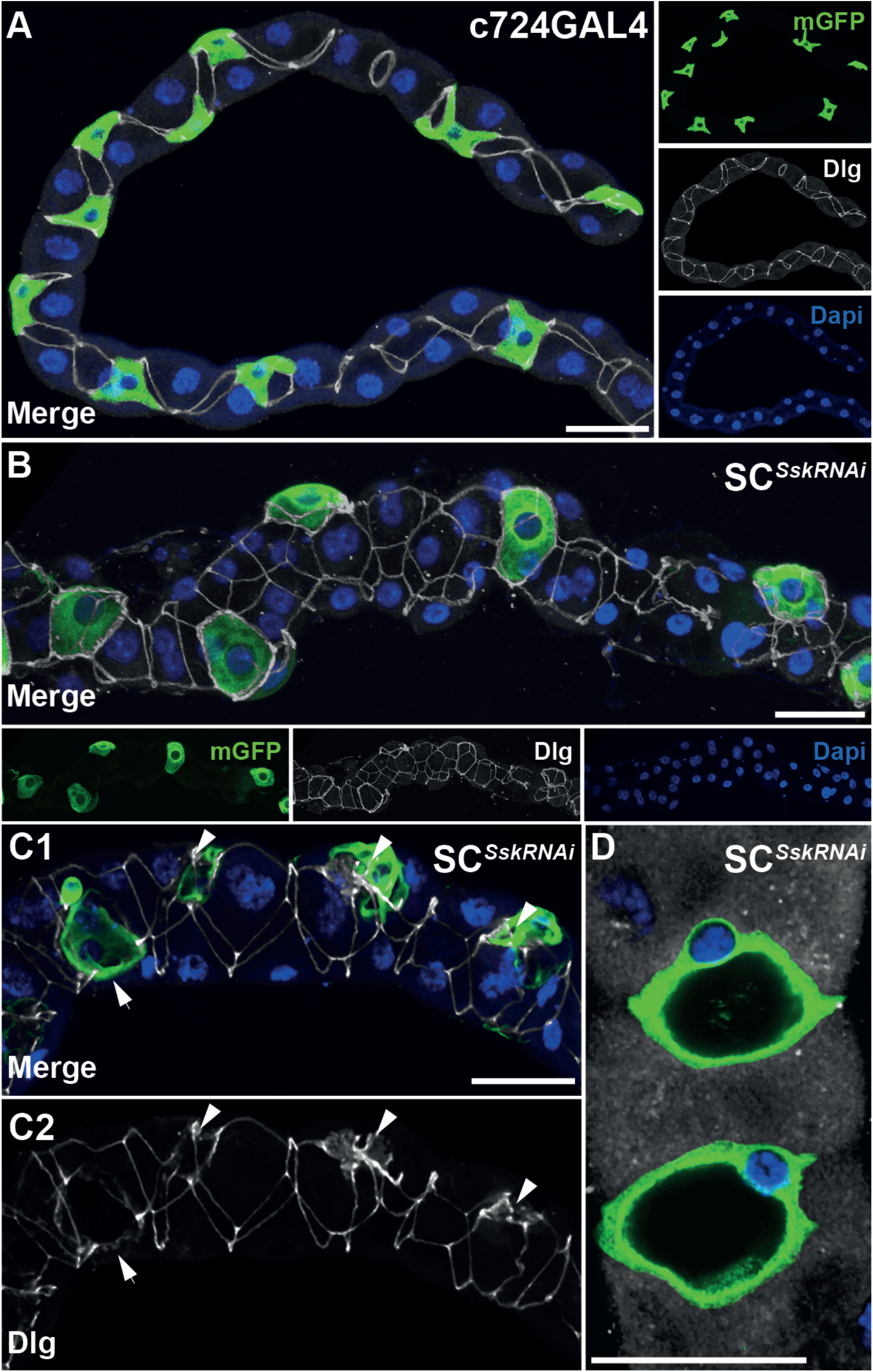
SC-specific *Ssk* depletion in adult Malpighian tubules results in compromised cellular morphology. (A) *c724*GAL4 driving membrane-bound (mGFP) in 5 day adult MT with expression in evenly spaced SCs exhibiting stereotypical stellar morphology, and smoothly organised junctions throughout realized by anti-discs large (Dlg). (B) 5 day adult SC^*Ssk*RNAi^ MT. SCs exhibiting absence of mature stellar morphology and extrude from plane of MT. Junctional complexes appear disorganised or missing. (C) 5 day adult SC^*Ssk*RNAi^ MT highlighting accretions of Dlg associated with mutant SCs (D2; arrowheads). (D) Subset of z-stack of SC^*Ssk*RNAi^ MT detailing ‘inflated’ vacuolar mutant SCs. mGFP, green; Dlg, white; Dapi, blue. Scale bars = 50 μm.

As it has been shown that dysregulation of Dlg results in disruption of micro-filament maintained cytoarchitecture (Izumi et al., 2019; Woods et al., 1996), we examined MTs in which *Ssk* had been knocked down and found F-actin micro-filaments were clearly absent in SC^*Ssk*RNAi^ SCs, while filaments associated with bicellular boundaries appeared reduced (Figure 3B). This loss of internal cytoarchitecture is almost certainly a contributing factor in the failure of SC^*Ssk*RNAi^ SCs to develop mature stellar morphology. This loss of cytoarchitecture was also apparent in PC^*Ssk*RNAi^ MTs (Figure S2C). Dysregulation of Dlg, and the associated loss of cytoarchitecure, also results in failure of apicobasal polarity (Laprise and Tepass, 2011). This is consistent with the observed loss of apical bias in GFP expression in experimental PC^*Ssk*RNAi^ MTs as compared with controls (Figure S2A-B). To further test this idea we employed an antibody to the Na^+^/K^+^ ATPase alpha-subunit – a transporter known to localize to the basolateral membrane (Patrick et al., 2006) – as an indicator of loss of apicobasal polarity. These data showed an overall reduction of Na^+^/K^+^ ATPase expression in SC^*Ssk*RNAi^ MTs as compared to controls, with a marked decrease in basal localisation in SCs indicative of a loss of apicobasal polarity (Figure S3). This loss is perhaps not surprising considering the functional requirement for proper formation of junctional complexes in determining overall cellular polarity (Bonello et al., 2019; Nelson, 2003), but importantly these phenotypes appear progressive, becoming more overt as experimental adults age.

**Figure 3.**
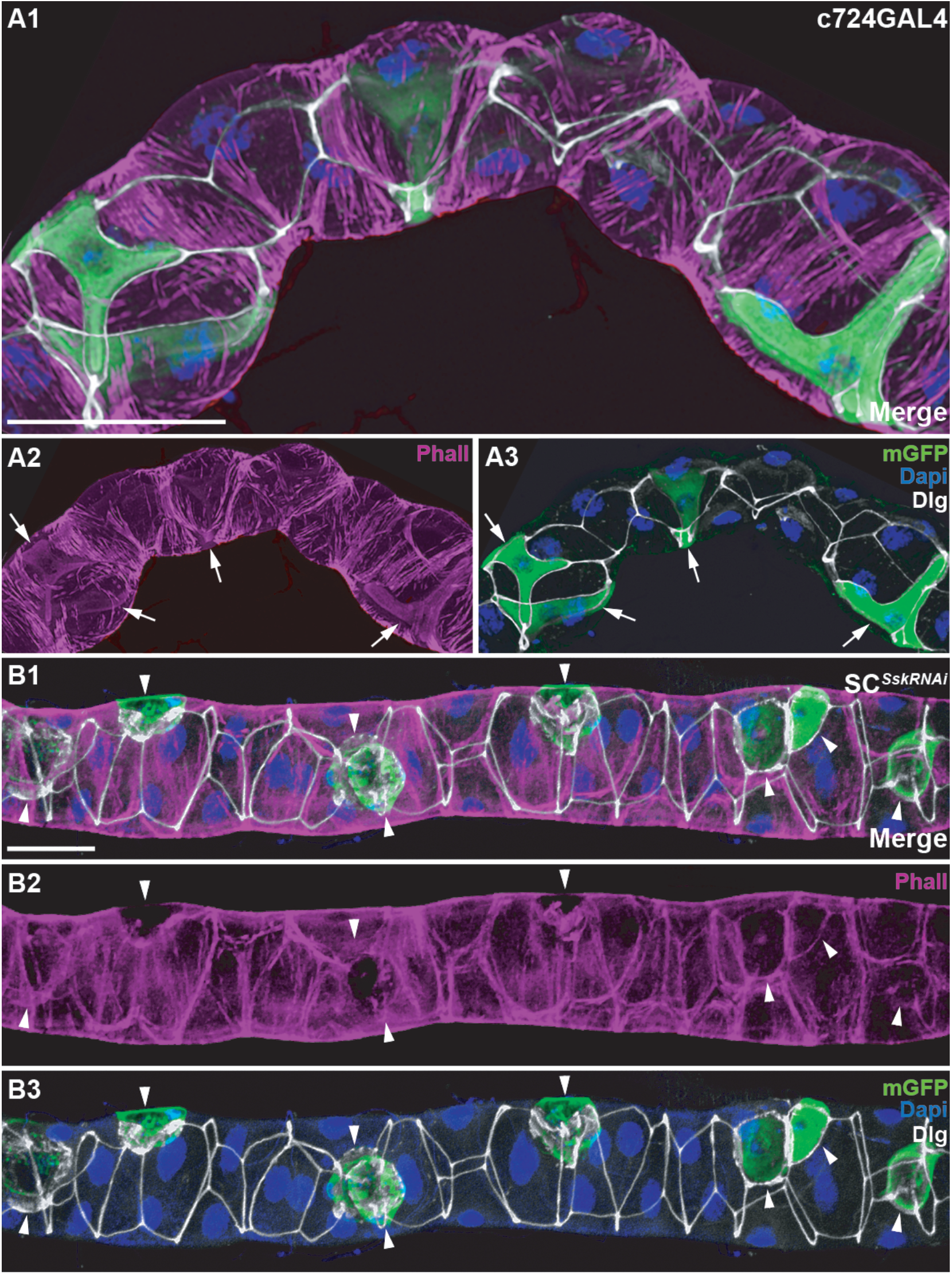
SC-specific depletion of *Ssk* in adult Malpighian tubules results in loss of cytoarchitectural organisation. (A) *c724*GAL4 driving membrane-bound (mGFP) in 5 day adult MT with cellular architecture, strongly associated with cellular junctions, realised by Phalloidin (Phall; F-actin) staining. A2-A3 highlighting internal cytoarchitecture specific to SCs (arrows). (B) 5 day adult SC^*Ssk*RNAi^ MT highlighting loss of F-actin associated with cellular junctions and SCs. B1-B2, highlighting absence of internal SC cytoarchitecture (arrowheads). Accretions of Dlg overt in degenerating SCs with reduced GFP expression and loss of Dapi staining. mGFP, green; Phall, magenta; Dlg, white; Dapi, blue. Scale bars = 50 μm.

In SCs, kinin-modulated Cl^−^ shunt conductance occurs specifically through the chloride channel ClC-a localizing to the basolateral membrane (Cabrero et al., 2014, 2020). We therefore employed anti-ClC-a to assay polarity in SC populations in experimental adult MTs and found ClC-a expression absent in the cuboidal SC population in SC^*Ssk*RNAi^ MTs (Figure S4B) but apparently unaffected in SCs in PC^*Ssk*RNAi^ MTs (Figure S4C). To test the physiological significance of these cellular defects, we exposed animals to conditions known to induce osmotic stress and assayed for organismal survival. These results revealed that SC^*Ssk*RNAi^ adults demonstrated a significant reduction in survival when allowed access to water only (non-desiccating starvation) (Figure S5A), but intriguingly this compromised viability is absent when exposed to high salt loading (Figure S5B). These data are consistent with the observed defects in SC function and compromised hormone-induced change in Cl^−^ and water fluxes in SC^*Ssk*RNAi^ animals (Cabrero et al, 2014; Denholm et al., 2013; Feingold et al, 2019), which result in a reduced capacity to respond and adapt to hypoosmotic challenges and regulate systemic fluid balance. Taken together, our data demonstrates a necessary role for *Ssk* in maintaining the SC transport machinery and by extension tubule transport competency and organismal fluid balance.

### Cell-specific depletion of Ssk results in absence of septa and impairment of junctional, cellular and tissue organisation

We next set out to determine the cellular mechanisms through which knock down of *Ssk* in specific sub-population of cells in the adult MTs could then affect overall tissue morphology and homeostatic capabilities. While both SC^*Ssk*RNAi^ and PC^*Ssk*RNAi^ MTs appear hyperplastic (Figures 2B and S2B), there is only a small, though significant, increase in overall population of cells in PC^*Ssk*RNAi^ MTs (Figure 4A). In contrast there is a slight decrease in the SC population in anterior SC^*Ssk*RNA^ MTs (Figure 4A-B), presumably due to progressive loss of SCs through apoptosis.

**Figure 4.**
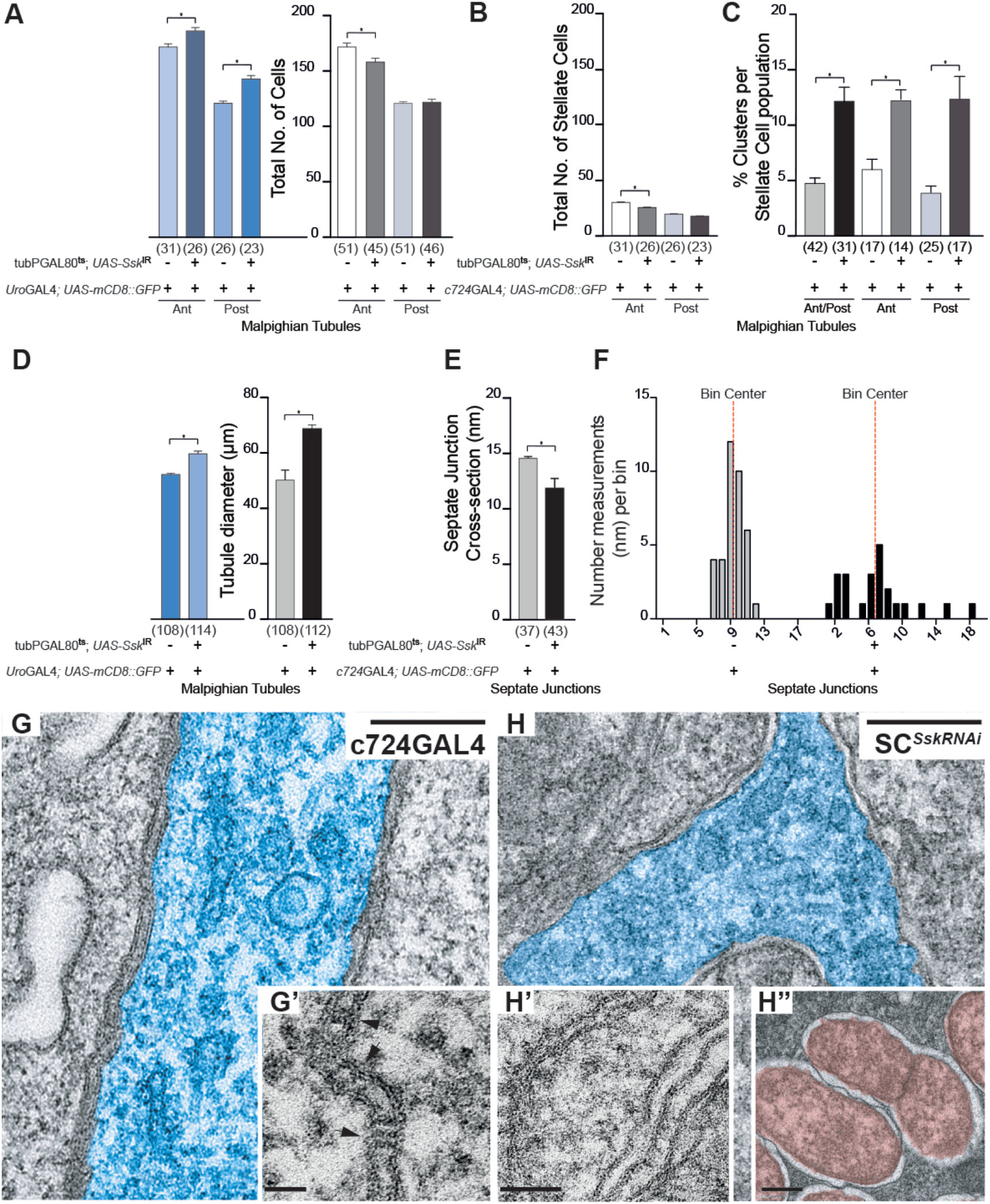
Cell-specific depletion of *Ssk* in adult Malpighian tubules results in failure of septa and junction, cell and tissue organisation. (A) Comparison of total cell counts for SC^*Ssk*RNAi^ and PC^*Ssk*RNAi^ MTs as compared to controls. Both anterior and posterior PC^*Ssk*RNAi^ MTs exhibit a significant increase in cell populations as compared with controls (186 ± 3, n=26 vs 171 ± 4, n=31, P<0.005 anterior tubules; 143 ± 4, n=23 vs 121 ± 2, n=26, P<0.005 posterior tubules). Anterior SC^*Ssk*RNAi^ MTs exhibit a significant decrease in cell populations while there are no changes in posterior tubules as compared with controls (158 ± 4, n=45 vs 174 ± 2, n=51; P<0.0001 anterior tubules; 122 ± 3, n=46 vs 121 ± 2, n=51; n/s posterior tubules). (B) Anterior SC^*Ssk*RNAi^ MTs exhibit a significant decrease in SC populations while there are no changes in posterior tubules as compared with controls (SC counts 25 ± 1, n=45 vs 30 ± 1, n=51; P<0.0002 anterior tubules; 18 ± 1, n=46 vs 19 ± 1, n=51; n/s posterior tubules). (C) Both anterior and posterior SC^*Ssk*RNAi^ MTs exhibit a significant increase in % SC cell clusters compared with controls (12.2 ± 1.27, n=31 vs 4.7 ± 0.59, n=42, P<0.0001). Cluster defined as two or more adjoining SCs. (D) Comparison of tubule cross sectional diameters for SC^*Ssk*RNAi^ and PC^*Ssk*RNAi^ MTs as compared to controls. Both anterior and posterior PC^*Ssk*RNAi^ and SC^*Ssk*RNAi^ MTs exhibit a significant increase in tubule cross sectional diameter as compared with controls (69.10 nm ± 1.184, n=114 vs 50.30 nm ± 3.883, n=108; P<0.0001 and 60.24 nm ± 0.8628, n=112 vs 52.75 nm ± 0.6783, n=108; P<0.0001 respectively). (E) SC^*Ssk*RNAi^ septate junctions exhibit a significant decrease in cross-sectional distance as compared with controls (11.77 nm ± 0.868, n=43 vs 14.43 nm ± 0.199, n=37; P<0.0001). (F) Junctional spans for SC^*Ssk*RNAi^ MTs are highly irregular, as evidenced by the frequency-distribution in septate junction measurements. N’s in parentheses. *P<0.0001. (G) TEM of 5 day old *c724*GAL4 control MT featuring the septal junctions associated with an SC (pseudocolour, blue), demonstrating the regular spacing and stereotypical ladder appearance caused by the presence of the septa. Scale bar = 0.2 μm. G’ inset, higher magnification image highlighting the presence of septa (arrows) spanning the septate junctions. Scale bar = 50 nm. (H) TEM of 5 day old SC^*Ssk*RNAi^ MT featuring the septal junctions associated with a SC (pseudocolour, blue), demonstrating disorganised spacing and absence of septa ‘ladders’. Scale bar = 0.2 μm. H’ inset, higher magnification image highlighting the absence of septa associated with mutant septate junctions. H” inset, higher magnification image highlighting the presence of dividing bacterium (psuedocolour, red). Scale bars = 50 nm.

In SC^*Ssk*RNAi^ SCs, not only is mature stellar morphology affected, but so too is spatial distribution; with a significant increase in the %SC population exhibiting clustering of 2 or more cells (Figure 4C). Tubules are traditionally regarded as developmentally ‘static’, that is the SC population has intercalated and been positioned within the tubule primordia by mid-embryogenesis and are thought merely to ‘mature’ physiologically (developing their characteristic stellar or bar morphology during final stages of pupariation) (Denholm, 2013; Denholm et al., 2013; 2003; Dow, 2012). This dysregulation in positioning of the SC population is indicative that the processes that determine, and maintain, the MTs cellular architecture may extend past embryogenesis in the fly’s development.

Both SC^*Ssk*RNAi^ and PC^*Ssk*RNAi^ MTs also have an increased cross-sectional diameter (Figure 4D), a phenotype iterating pathological changes observed during intestinal barrier dysfunction (Izumi et al., 2019; Rera et al., 2013a; Salazar et al., 2018) and when Mesh or Tsp2A are impaired in MTs (Beyanbach et al., 2020; Jonusaite et al., 2020). Complementing these observations, while TEM scans show the septal junction span is significantly reduced (Figure 4E), the junctional spans are in fact highly irregular (as evidenced by the frequency-distribution in septal junction measurements; Figure 4F). Importantly there is a complete absence of septa (Figure 4H and H’), a consequence of which is loss of para-cellular barrier function (Baumgartner et al., 1996; Genova and Fehon, 2003; Lamb et al., 1998). Absence of septa does not itself result in loss of septal-gap structural integrity, as demonstrated by septal mutants such as *sinuous* and *coracle* (Lamb et al., 1998; Wu et al., 2004). Rather, structural integrity is a function of the adherens junctions (AJ) (Tepass and Hartenstein, 1994). Dlg localisation however has been shown to be regulated by, and in turn modulate, AJ formation (Bilder et al., 2000; 2003; Bonello, et al., 2019; Harris and Peifer, 2004), so that failure of AJ’s may be an effect of the mislocalisation of Dlg occurring as a consequence of Ssk impairment, which then manifests in the failure of septal-gap structural integrity.

Loss of septa results in failure of paracellular barriers, allowing opportunistic toxic and/or pathogenic invasion (Izumi and Furuse, 2014; Salazar et al., 2018). TEMs of experimental animals show evidence of bacterial invasion not observed in controls (Figure 5H’’). This increasing bacterial load would result in chronic activation of inflammatory/immune responses (Izumi et al., 2019; Rera et al., 2013a; 2012; Resnick-Docampo et al., 2018; Salazar et al., 2018) again iterating pathological phenotypes observed during intestinal barrier dysfunction, contributing to reduced organismal viability (Izumi et al., 2019; Rera et al., 2013a; Salazar et al., 2018).

**Figure 5.**
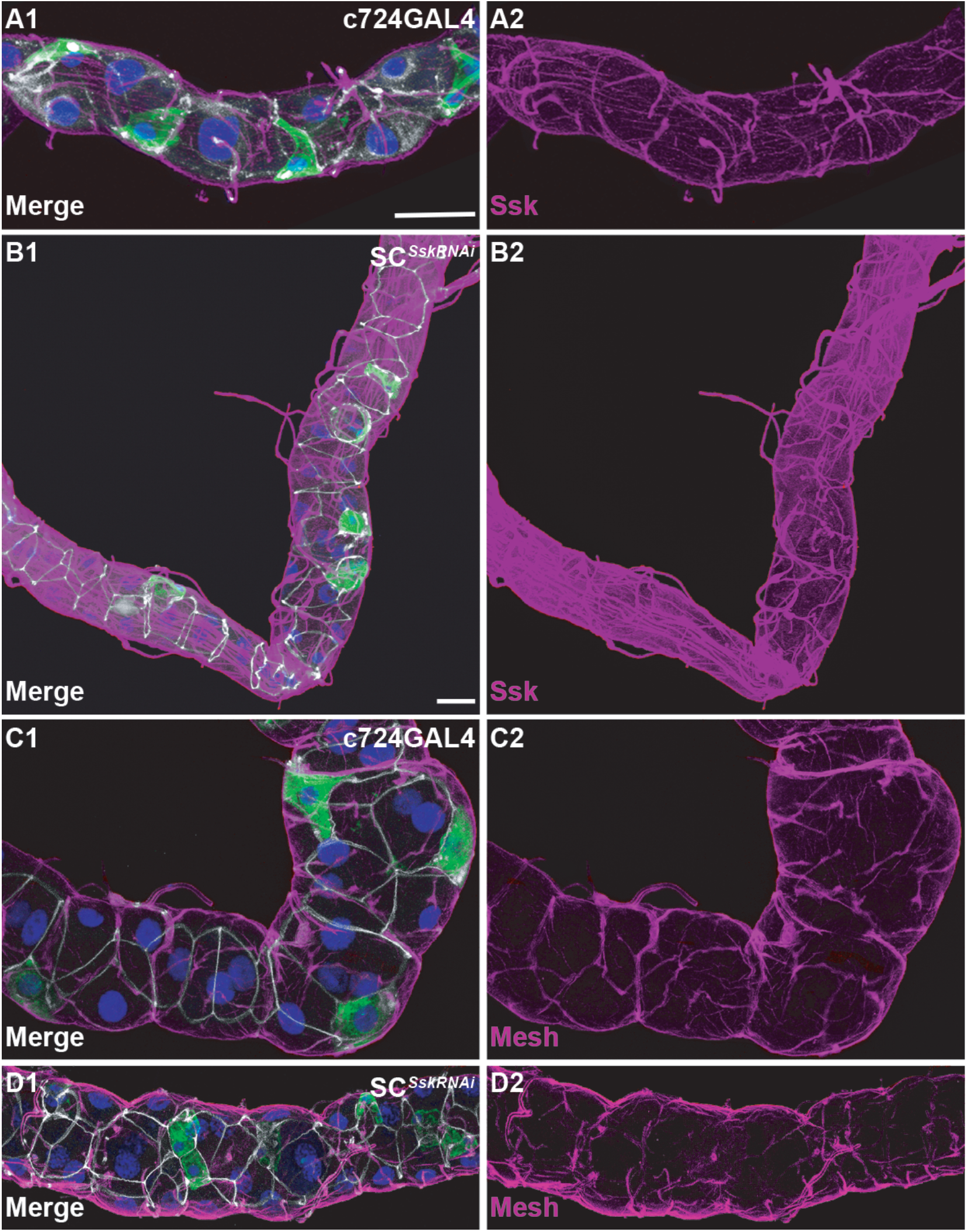
Cell-specific depletion of *Ssk* results in hyperplasia of trachea supplying adult Malpighian tubules. (A) *c724*GAL4 driving membrane-bound (mGFP) demonstrating control levels of Ssk expression associated with junctions and (at lower levels) trachea in a 5 day adult MT. (B) Overt hyperplasia of trachea, realised by Ssk expression, associated with main segment of SC^*Ssk*RNAi^ 5 day adult MT. (C) *c724*GAL4 driving mGFP in a 5 D adult MT, demonstrating control levels of Mesh expression associated with junctions and (at lower levels) trachea. (D) SC^*Ssk*RNAi^ 5 D adult MT demonstrating a disorganised and reduced pattern of Mesh expression. Nb-control and experimental confocal stacks for (A – B) and (B – C) were collected using identical settings. mGFP, green; Ssk or Mesh, magenta; Dlg, white; Dapi, blue. Scale bars = 50 μm.

Taken together these results demonstrate that cell-specific depletion of *Ssk* not only compromises individual cellular junctional complexes but affects tubule morphology and organisation as a whole, compromising barrier integrity and homeostatic capabilities and allowing opportunistic pathogenic invasion, all of which contributes to significantly reduced organismal viability. That our findings show impairment of junctional complexes in a restricted population of cells affects overall tissue function must also speak to the idea that these cell-cell ‘tight’ junctions are required to direct appropriate non-cell autonomous communication(s).

### Cell-specific Ssk depletion results in proliferation of tiny cells and hyperplasia of trachea supplying the MTs

We assessed the efficacy of the *Ssk*^RNAi^ transgene via qPCR analysis and immunolocalisation using an antibody specific to Ssk. We were able to confirm a tissue-specific knockdown of Ssk of ~35% in 5 day old adult SC^*Ssk*RNAi^ MTs as compared with controls (Fig. S6A). However, immunolocalization of the Ssk Ab yielded the surprising observation that not only do the trachea supplying the MTs also express Ssk, but that there is an overt hyperplasia of this trachea in 5 day old SC^*Ssk*RNAi^ MTs (Figures 5B). So the actual level of Ssk knockdown within the experimental MTs is likely masked by the increased expression of Ssk in the closely associated hyperplastic trachea. This hyperplasia of the trachea supplying the MTs is also apparent in PC^*Ssk*RNAi^ MTs, though not to the levels observed with SC^*Ssk*RNAi^ MTs (Figure S6C). Application of an antibody specific to Mesh, *Ssk*’s obligate partner (Izumi et al., 2012), demonstrated that distribution of Mesh at the junctional complexes appeared reduced and disorganised in SC^*Ssk*RNAi^ MTs, but did not manifest the same increased expression in hyperplastic trachea as observed with Ssk (Figure 5D).

In both SC^*Ssk*RNAi^ and PC^*Ssk*RNAi^ MTs we also observed an increased number of nuclei associated with ‘tiny cells’ in the proximal area of the tubule and ureter (Figures S7). These tiny cells are described as renal and nephritic stem cells (Martinez-Corrales et al, 2019; Singh and Hou, 2008; 2009; Singh et al., 2011; Wang and Spradling, 2020) and are observed to increase in number in response to stress or as MTs degenerate over time. In keeping with these observations, some of these cells co-express the proliferative cell marker Delta (Biteau et al., 2008) (Figure S7C-D). However, we cannot rule out that this may be a secondary effect of epithelial dysfunction causing anomalous expression of proliferative markers as has been observed to occur during gut barrier dysfunction when some intestinal enterocytes express ISC proliferative markers (Resnik-Docampo et al., 2017).

It is intriguing that the observed proliferation of (potential) stem cells and hyperplasia of associated trachea is coincident with mislocalisation of the cell growth regulator Dlg (Bonello et al., 2019; Elsum et al., 2012; Humbert et al., 2008; 2003; Khoury et al., 2020). Whether these effects are causally related or merely coincidental, reflecting overall failure in tissue homeostatic functions due to dysregulation of the junctional complexes, warrants further investigation.

### Ssk impairment accelerates age-related changes in tubule septate junction integrity and secretory capacity

To assess potential age-related changes in tubule physiology, we examined junctional integrity by analysing Dlg localization as well as renal secretory capacity, at progressive time-points during ageing. We observed a significant decrease in Dlg intensity associated with cell-cell junctions in older as compared with younger animals (Figure 6A-B), as well as a progressive decline in secretory activity, as evidenced by significant reductions in both basal and stimulated rates of tubule secretion (Figure 7A). These data suggest that junctional stability declines with age and that there is a measurable ‘natural’ deterioration in transport capacity of the tubule that correlates with physiological ageing (Figure 7A-B).

**Figure 6.**
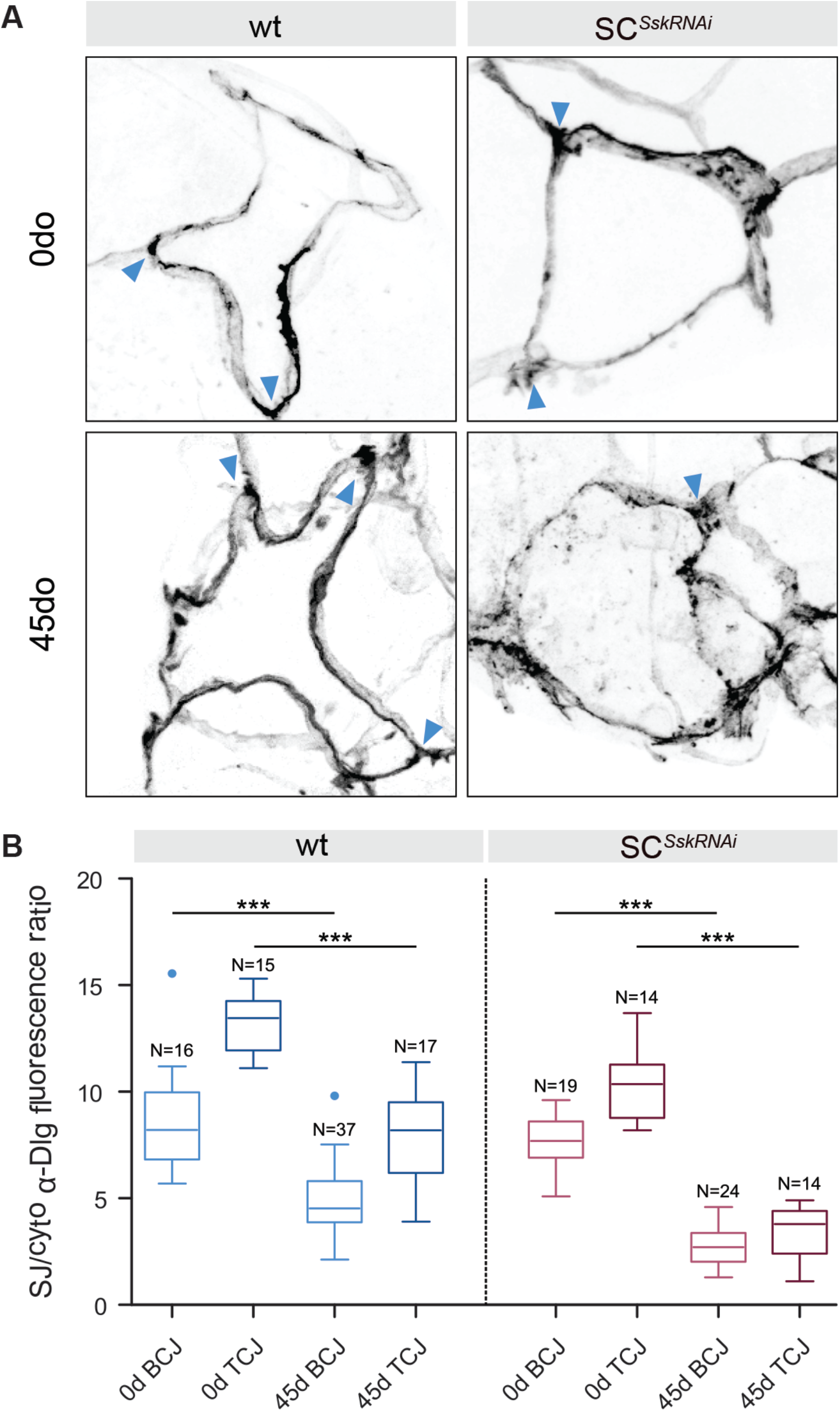
SC-specific depletion of *Ssk* impairment advances age-related changes in junctional protein localisation. (A) Distribution of Dlg in adult wt and SC^*Ssk*RNAi^ SCs at 0 and 45 day old (post-eclosion) time periods. tricellular junctions indicated by arrowheads. (B) Graphical representation of the ratio of Dlg fluorescent signal localised to the bicellular (BCJ) and tricellular (TCJ) junctions compared with cytoplasm in adult wt and SC^*Ssk*RNAi^ SCs at 0 and 45 day old (post-eclosion) time periods.

**Figure 7.**
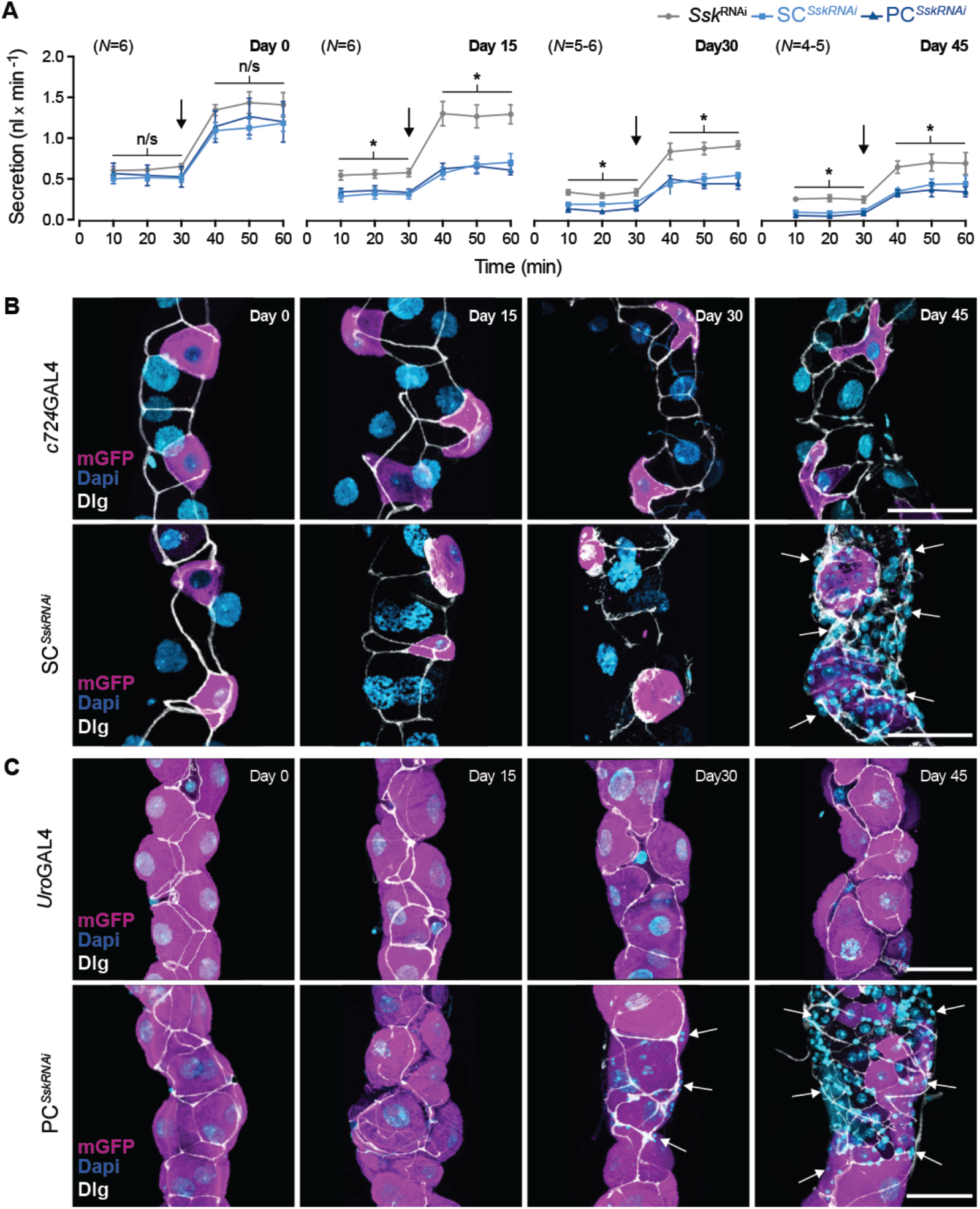
*Ssk* impairment accelerates age-related changes in septate junction integrity and secretory capacity in adult Malpighian tubules. (A) Modified Ramsey assay measuring fluid secretion rates in control and experimental adult MTs over time as flies age. The secretion rate capacity for all tubules (both control and experimental) decreases progressively over time. However, secretion rates post-day 15 for SC^*Ssk*RNAi^ and PC^*Ssk*RNAi^ MTs are significantly reduced as compared to controls, even after stimulation with Dromekinin (black arrows). *P<0.05, paired samples t-test. N’s in parentheses. Realisation of the accelerated degeneration of tubule morphology, mislocalisation of Dlg in SC^*Ssk*RNAi^ (B) and PC^*Ssk*RNAi^ (C) MTs as compared with controls. Post-day 30 proliferation of tiny cells in SC^*Ssk*RNAi^ and PC^*Ssk*RNAi^ MTs overt (arrows). mGFP, magenta; Dlg, white; Dapi, blue. Scale bars = 50 μm.

We also investigated whether compromising junctional integrity, through cell-specific depletion of *Ssk*, could phenocopy these age-related changes in tubule function. Knocking down Ssk expression, in either SC^*Ssk*RNAi^ or PC^*Ssk*RNAi^ MTs, resulted in an accelerated and progressive decline in tubule secretory capacity relative to age-matched controls (Figures 7A and S8). This decline is evident as early as 15 days post-eclosion, with this decrement in function mirrored by an associated degeneration in tissue morphology; with overt signs of apoptosis, mislocalised cytoplasmic Dlg accretions and a dramatic increase in ‘tiny’ cells (Figures 6A-B, 7A-B). Compellingly, when comparing secretory rates between 45 day old control and 15 day old experimental animals, while the experimental animals’ basal secretory rate were marginally better, there was no significant difference in stimulated rates of tubule secretion between control and experimental animals (Figure S8). That is, the 15 day old experimentally aged tubule recapitulated the reduced secretory capacity of 45 day old controls that was a result of ‘natural’ physiological ageing. That compromised junctional integrity resulted in an accelerated loss in tissue homeostasis is supported by the observed significant increase in Dlg mislocalisation in SC^*Ssk*RNAi^ MT cellular junctions as compared with controls of the same age (Figures 6A-B).

Our investigations demonstrate, for the first time, a measurable age-dependent decline in secretory transport capacity in adult tubules, that correlates with mislocalisation of SJ proteins and coincident progressive degeneration in cellular and tissue morphology. Critically, by cell-specific depletion of *Ssk*, we were able to phenocopy the failure of junctional complexes, leading to an accelerated reduction in secretory capacity. Providing insight into the mechanism by which failure in barrier integrity can advance loss of systemic fluid homeostasis, morbidity and, ultimately, death.

## DISCUSSION

Transporting epithelia must be able to proliferate appropriately in response to growth directives (organogenesis, wound response etc.), while acting to ensure cellular compartmentalization to ensure key systems are protected from physiological and xenobiotic stresses and provide a physical barrier against invasive pathogens, all the while allowing appropriate transport of ions, solutes and water. Pivotal to these diverse functional demands is the formation, and maintenance, of ‘tight’ cell-cell junctional complexes. The profound consequences associated with a failure in these processes have been demonstrated in intestinal epithelia dysfunction, resulting in physiological decline contributing to onset of age-related disease and, ultimately, death (Clark and Walker, 2018; Clark et al., 2015; Davies et al., 2012; Hu and Jasper, 2017; McGee et al., 2011; Regan et al., 2016; Rera et al., 2013a; 2012; Resnik-Docampo et al., 2017; 2018; Salazar et al., 2018).

### Ssk depletion phenocopies natural age-related changes in tubule junctional integrity affecting organismal viability

Our investigations demonstrate that a measurable ‘natural’ decline in tubule transport capacity, which correlates with physiological ageing, occurs over time. This decrement in tubule function occurs in conjunction with mislocalisation of cellular junctional components, indicative of a failure in cell-cell junctional integrity, resulting in a progressive decline in organismal homeostatic capabilities and viability. This is supported by Salazar et al., 2018, who demonstrated that inflammatory response genes are naturally upregulated as flies age, and that intestinal barrier function was equally compromised when comparing controls at day 45 with experimental flies in which Intestinal barrier dysfunction had been induced. Finally, we show that by restrictively impairing expression of the essential junctional protein Ssk (Yanagihashi et al., 2012), in either SCs or PCs, we can both phenocopy and advance normal degenerative changes, affecting not only individual cell populations but overall tissue morphology, resulting in a measurable accelerated decline in the tubule’s physiological capacities, and culminating in significantly reduced organismal viability.

### Ssk depletion promotes loss of cytoarchitecture and apicobasal polarity

Ssk, Tsp2a, and the membrane spanning protein Mesh are mutually dependent for their localisation to, and critical for, proper formation of the sSJ’s (Beyanbach et al., 2020; Jonusaite et al., 2020; Izumi et al., 2012; Izumi et al., 2016; Salazar et al., 2018; Xu et al., 2019; Yanagashi et al., 2012). This Mesh/Ssk/Tsp2a complex creates a platform for appropriate assembly and localisation of other key junctional proteins (Beyanbach et al., 2020; Jonusaite et al., 2020; Izumi et al., 2012; Izumi et al., 2016; Salazar et al., 2018; Xu et al., 2019). In pSJs it has been shown that Dlg is not a core junctional component, though its misregulation still dramatically affects junctional complexes (Oshima and Fehon, 2011; Izumi et al., 2012). In keeping with this observation, Mesh/Ssk/Tsp2a do not appear to directly interact with Dlg in sSJs, as their ability to localise to junctions are independent (Yanagashi et al., 2012; Izumi et al., 2012; Izumi et al., 2016). However, the individual components of the Mesh/Ssk/Tsp2a complex are essential to appropriate localisation of Lgl (Jonusaite et al., 2020; Izumi et al., 2012; 2016; Yanagashi et al., 2012), and there exists a mutually dependent functional relationship between Lgl and Dlg (Bilder et al., 2000; 2003; Izumi et al., 2012; Su et al., 2012). Dlg and Lgl, along with Scribble, comprise the ‘Scribble polarity module’, which acts as a regulator of cell polarity and proliferation (Bonello et al., 2019; Elsum et al., 2012; Humbert et al., 2008; 2003; Khoury et al., 2020). We propose that in the absence of Ssk, Lgl is mislocalised at the MTs’ apical membrane, which in turn causes mislocalisation of Dlg, disrupting the Scribble polarity module as well as affecting other components of the sSJs (*cf* Beyanbach et al., 2020; Jonusaite et al., 2020; Izumi et al., 2019). This mis-regulation of Dlg, evidenced by progressive formation of cytoplasmic accretions, results in disruption of micro-filament networks responsible for the internal cellular cytoarchitecture (Bilder and Perrimon, 2000; Bilder et al., 2003; Tanentzapf & Tepass, 2003; Woods et al., 1996; Wu & Beitel, 2004; Yu and Fernandez-Gonzalez, 2016), most evident in the dysmorphic SCs where F-actin appears entirely absent.

In SC^*Ssk*RNAi^ animals this mislocalisation of Dlg results in loss of apicobasal polarity (Bilder and Perrimon, 2000; Bilder et al., 2003; Bonello et al., 2019; Tanentzapf & Tepass, 2003; Woods et al., 1996; Wu & Beitel, 2004) as evidenced by the absence of basolateral expression of Na^+^/K^+^ ATPase. That cellular function and polarity is compromised is further demonstrated by the absence of the SC-specific chloride channel (ClC-a), normally present at the basolateral membrane (Cabrero et al., 2014), in the SCs of SC^*Ssk*RNAi^ animals. The absence of Clc-a in the SCs also means that the Cl^−^ shunt, necessary to create the osmotic gradient required for water flux (Cabrero et al., 2020; 2014; Denholm et al., 2013), is compromised, which again speaks to the decrement in osmoregulatory capacity observed in these tubules.

Impairment of polarity cues and cell growth regulators is a necessary presage for epithelium-mesenchymal transition in developing cancers (Elsum et al., 2012; Humbert et al., 2008; 2003; Royer and Lu, 2011; Wodarz and Näthke, 2007). It is of note that, during development, the SC cell population migrates and intercalates into the tubule primordia, undergoing a programmed mesenchymal-epithelium transition (Denholm et al., 2003). It would be tempting to speculate that in SC^*Ssk*RNAi^ MTs, should the mutant SCs not undergo premature apoptosis, this loss of polarity, in conjunction with failure to develop a mature stellar morphology, presentation of an extruded cellular profile and breakdown in cell-to-cell communication would indicate a shift towards development of further tumorigenic characteristics.

### Failure of septa and junctional structure results in compromised tubule barrier integrity

The MTs’ compromised fluid integrity is due to a loss of septa and therefore para-cellular barrier function. While reduction in viability in experimental animals must, initially at least, be a consequence of the significant progressive decline in osmoregulatory capacity resulting in physiological failures, this effect is compounded by loss of barrier functions, which allows opportunistic invasion of pathogens and has been shown to lead to dysbiosis and activation of immune/inflammatory responses (Clark and Walker, 2018; Clark et al., 2015; Izumi and Furuse, 2014; Rera et al., 2013b; Resnick-Docampo et al., 2018; Salazar et al., 2018; Xu et al., 2019). These effects phenocopying those described for age-related loss of intestinal integrity (Rera et al., 2012; Resnik-Docampo et al., 2017; 2020; Salazar et al., 2019).

Concomitant with loss of septa was an observed irregularity of the junctional spans, indicating a loss in junctional structural integrity, likely due to impairment of AJs. The AJs also provide cues for appropriate localisation of, and may then be modulated by, Dlg in junctional complexes (Bilder et al., 2000; 2003; Bonello et al., 2019; Harris and Peifer, 2004). AJs are also required during the integration of the SCs in the developing tubule, for proper localisation and polarity of the SC population during formation of the junctional complexes (Campbell et al., 2010). Therefore, any compromise of these structures may again contribute to the observed atypical SC distribution, loss of polarity and failure of the junctional complexes. Further, the fact that increased clustering of SCs occurs when *Ssk* expression is knocked down in pupal and adult stages might indicate that the MTs are not completely developmentally static and are able to respond to environmental cues in order to assure tissue homeostasis throughout the animal’s life.

### Cell-specific depletion of Ssk in tubule epithelium has global consequences

Impairment of the sSJ junctional complexes, with consequent loss of cell cytoarchitecture and polarity, has profound effects on the individual cells. However, these effects are not limited to the cells in which Ssk expression has been specifically impaired, but rather affects morphology and function across the tubule, with this degradation of epithelium competence affecting viability. That impairment of junctional complexes in a specific sub-population of cells may then affect overall tissue function and morphology speaks to the idea that non-cell autonomous communication(s) are required across these ‘tight’ junctions. This is supported by the observations that mutant gut clonal cells in which *Tsp2A* expression is impaired induced non-cell autonomous stem cell proliferation (Izumi et al., 2019) and that *mesh* knock-down in PCs resulted in dysmorphic SC development (Jonusaite et al., 2020).

Realisation of the apparent dysregulation of Dlg, a known tumour-suppressor gene (Elsum et al., 2012; Humbert et al., 2008; 2003), in individual junctional complexes may also speak to the effects that occur more universally throughout the mutant MTs. Polarity proteins have been shown to directly modulate signalling pathways controlling tissue growth (Royer and Lu, 2011; Tepass et al., 2001) and certainly one of the most striking global effects observed in both SC^*Ssk*RNAi^ and PC^*Ssk*RNAi^ MTs is the proliferation of ‘tiny’ cells at the ureters and the hyperplasia that occurs in the associated trachea supplying the MTs. Whether the trachea hyperplasia is an indirect effect of local tissue hypoxia due to a compensatory upregulation of metabolic function, as seen during fly development (Texada et al., 2019), or is a direct consequence of impairment of appropriate tissue growth regulation due to dysregulation of the junctional complexes warrants further investigation. However, it is clear that the progressive proliferation of the tiny cells, which have been described as potential renal and nephritic stem cells (Martinez-Corrales et al., 2019; Singh and Hou, 2008, 2009; Singh et al, 2011; Wang and Spradling, 2020), occurs in direct response to the failing of the tubules’ physiological capacity. Again further investigation is needed to establish if these cells are actually proliferative in nature, or if they are anomalously expressing proliferative markers, as was observed with enterocyte-like cells expressing ISC proliferative markers during intestinal barrier dysfunction (Resnik-Docampo et al., 2017). However it is not unreasonable to relate these effects, occurring in conjunction with, or as result of, loss-of-barrier function allowing opportunistic pathogenic challenge and consequent dysbiosis, to the inflammatory and ageing responses involving the JNK and Jak/Stat/Cytokine signalling paths that direct increased proliferation of stem cells and other regenerative homeostatic processes in the gut (Biteau et al., 2008; Izumi et al., 2019; Jiang et al., 2009; Rera et al., 2012; Resnik-Docampo et al., 2017; 2020).

Altogether, our findings describe an essential role for *Snakeskin* in appropriate formation of smooth septate junctions, ensuring epithelial integrity and acting, directly or indirectly, as a modulator for tissue growth in *Drosophila* Malpighian tubules. We demonstrate that a significant and progressive decline in secretory transport capacity in the adult MTs occurs naturally over time as result of, or in conjunction with, a failure of cellular junctional components, with the overall tissue displaying progressive degeneration in cellular morphology. We demonstrate that cell-specific impairment of *Ssk* in the adult MTs accelerates the mislocalisation of junctional components and degenerative effects in tubule morphology, with an associated measurable decline in functional capacity, resulting in significantly reduced viability. These investigations highlight the pivotal role that cell-cell junctional integrity plays in assuring epithelial competence, providing insight into how age-related progressive degeneration of junctional components can lead to failure of cellular, tissue and organismal homeostasis resulting in advancing morbidity and, ultimately, death.

## Acknowledgements

We would like to thank Prof. Mikio Furuse for the kind supply of fly lines and reagents. We would also like to thank Dr. Selim Terhzaz for invaluable help with qPCR analyses and critical comments. Thanks to Margaret Mullin of the Electron Microscopy Facility, School of Life Sciences, MVLS for her assistance and to Prof. Stephen Goodwin, Dr. Adam Dobson, Mr. Pablo Cabrero and the Dow Davies lab for critical comments and advice. This work was supported by funding from UKRI BBSRC (BB/P008097/1) to SAD and JATD with additional funding given by the Villum Foundation (15365) to KAH.

## MATERIALS AND METHODS

### Drosophila stocks

*Drosophila* lines were reared on standard Glasgow *Drosophila* diet at 45-55% relative humidity with a 12:12 hour light:dark photoperiod at a temperature of 22°C unless otherwise stated. Parental (control) and F1 progeny (experimental) strains expressing the *tubPGAL80*^*ts*^ transgene were raised at the permissive (18°C) temperature before the F1 (experimental) progeny were transferred at late L3/white pre-pupal stage to the restrictive (29°C) temperature*. w*; tubP GAL80*^*ts*^; *TM2/TM6b Tb*^*1*^ (7108) and *y*^*1*^ *w*; Pin*^*Yt*^/*Cyo*; UAS-*mCD8::GFP* (5130) lines acquired from Bloomington Stock Center (Bloomington, IN, USA). UAS-*Ssk*^RNAi^ line (Yanagihashi et al., 2012), kind gift of the Furuse Lab. The *c724*GAL4 (Sözen et al., 1997) and *Uro*GAL4 (Terhzaz et al., 2010) lines were generated in-house previously.

| Parental lines used to generate SC<sup>SskRNAi</sup> and PC<sup>SskRNAi</sup> experimental lines. |  |
| --- | --- |
|  | *w<sup>\*</sup>*; *tubPGAL80<sup>ts</sup>*; *UAS-Ssk<sup>RNAi</sup>* |
| SC-specific GAL4 driver | *w<sup>\*</sup>*; *c724GAL4*; *UAS-mCD8::GFP* |
| PC-specific GAL4 driver | *w<sup>\*</sup>*; *UroGAL4*; *UAS-mCD8::GFP* |

### Immunocytochemistry

Unless stated adult flies were reared at 29°C and aged for 5 days prior to dissection in Schneider’s *Drosophila* medium (Gibco, Thermo Fisher Scientific). Immunocytochemistry staining as previously described (Halberg et al., 2015). Primary antibodies employed, mouse monoclonal anti-A5 Na^+^/K^+^ vATPase α-subunit (1:50), −43F discs-large 1 (1:500), -C594.9B Delta (1:200) all DSHB (Univ. of Iowa, IA, USA); rabbit polyclonal anti-Snakeskin and -Mesh (1:1000), Furuse Lab (Izumi et al., 2012; Yanagihashi et al., 2012); and rabbit polyclonal anti-Clc-A (1:50) (Cabrero et al., 2014). Secondary antibodies, anti-rabbit Alexafluor 488 or 546 or anti-mouse Alexafluor 633 (Thermo Fisher Scientific) (1:600). Tubules were also incubated in 500 ng/ml DAPI and Phalloidin TRITC (Sigma-Aldrich). Tissues were mounted on either Polysine slides (VWR International, Leuven) or glass bottomed dishes (MatTek Corporation, MA, USA) and analysed using Zeiss a LSM 880 confocal micro-system (Carl Zeiss Ltd., Cambridge UK). Confocal Z projection stacks used for cell counts, analyses, and presentation were opened in ImageJ (NIH, MA, USA) prior to transferal to Adobe Photoshop and Illustrator (CS6; CA, USA) for final presentation.

### Transmission Electron Microscopy

Tubules were dissected from control and experimental adult flies and fixed in trialdehyde consisting of: 2.5% glutaraldehyde, 2% paraformaldehyde, 1.25% acrolein and 2.6% DMSO in 0.1M sodium cacodylate buffer (pH 7.4) overnight at 4°C. The tissues were post fixed in 1% OsO_4_ buffered with 0.1 M sodium cacodylate adjusted to pH 7.4 for 1 hour at room temperature. Next, the tubules were dehydrated through graded series of ethanol and propylene oxide, before being embedded in epoxy resin EPON 812 (TAAB, Berkshire, UK). Ultrathin sections were cut with a diamond knife on a Leica UTC ultramicrotome and stained with half-saturated (2%) uranyl acetate followed by Reynolds’ lead citrate. The ultrathin sections were examined in a FEI Tecnai T20 electron microscope.

### RNA isolation, cDNA synthesis and quantitative (q)-RT-PCR

RNA was isolated from 30 pairs of dissected adult tubules using TRIzol Reagent (Thermo Fisher Scientific, Renfrew, UK) resuspended in nuclease-free dH2O. cDNA was synthesized from 500 ng RNA using SuperScript II RT (Thermo Fisher Scientific, Renfrew, UK), following manufacturer’s instructions. q-RT-PCR was performed using Brilliant III SYBR Green QPCR Master Mix (Agilent) on the StepOne+ Real-Time PCR system (Thermo Fisher Scientific, Renfrew, UK) using the following primers specific to *snakeskin*, SskF1 - TTACACTGGATGCCACACCATTGC, SskR1 - TGACGCTCCGAGTTCACATACAGG, and *alpha tubulin 84b* TubF1-*CCTCGAAATCGTAGCTCTACAC*, TubR1 -*ACCAGCCTGACCAACATG* [Ref] (Integrated DNA Technologies). Following amplification, StepOne software was used to generate a standard curve. Relative concentration was determined by placing the Cycle Threshold (Ct) value and the values from the gene standard onto the standard curve. Each sample was then normalized against alpha-tubulin, resulting in a ratio of gene/alpha-tubulin expression. Results were then plotted as mRNA mean amounts ± s.e.m. using Prism 6.0 (GraphPad, CA, USA).

### Fly Weight Measurements

To measure wet-body weight, 20 female flies (n=8-12 groups) were anesthetized, transferred to pre-weighed Eppendorf tubes and weighed. For dry-body weight, flies were killed by freezing for 20 min and then dried at 60°C for 48 hr. Dry flies were weighed after reaching room temperature. All weighings were performed on a GR-202 analytical balance (A&D Instruments Ltd., Abingdon, UK).

### Survival assays

All survival assays comprised a minimum of 180 flies (n= 30 flies per vial, 3 vials per sex) and were performed in 12:12 hr LD at the restrictive (29 °C) temperature. Starvation assay vials contained 7 ml of 1% LMP-Agarose in water. Sucrose assay vials contained 7 ml of 1% LMP-Agarose in water with addition of 5% sucrose. Osmotic stress assays comprised vials containing either 7 ml of standard fly medium with the addition of 3% NaCl or with the addition of 1% H_2_O_2_. The age of the flies ranged between 3 – 5 days. Data for all assays were plotted as Kaplan–Meier curves and analysed using the Mantel-Cox (Log-rank) test using Prism v6.0 (GraphPad, CA, USA).

### >Ramsay fluid secretion assay

Fluid secretion assays using *Drosophila* Malpighian tubules were performed as described previously (Dow et al., 1994). Malpighian tubules were dissected in ice cold Schneider’s medium and transferred to a 9 μl drop of 1:1 of Schneider’s medium and *Drosophila* saline. Baseline secretion was measured every 10 minutes for 30 minutes, after which 1 μl of DromeKinin peptide (DK; 10^−6^ M) (Terhzaz et al., 1999) was added to the drop and stimulated secretion was measured every 10 minutes for a further 30 minutes. Percentage change of basal secretion rates were calculated as previously shown (Dow et al., 1994).

## Author contributions

AJD, KAH and JATD designed and conceptualized the study with input from SAD. AJD, KAH and LKB performed the experiments. AJD and KAH analyzed the data, wrote the manuscript and produced the figures. All authors reviewed and edited the manuscript.

## References

Baumgartner, S., Littleton, J.T., Broadie, K., Bhat, M.A., Harbecke, R., Lengyel, J.A., Chiquet-Ehrismann, R., Prokop, A., and Bellen, H.J. (1996). A Drosophila neurexin is required for septate junction and blood-nerve barrier formation and function. Cell 87, 1059–1068.

Beyenbach, K.W., Schoene, F., Breitsprecher, L.F., Tiburcy, F., Furuse, M., Izumi, Y., Meyer, H., Jonusaite, S., Rodan, A.R., and Paululat, A. (2020). The Septate Junction Protein Tetraspanin 2A is critical to the Structure and Function of Malpighian tubules in Drosophila melanogaster. Am J Physiol, Cell Physiol 141, 899.

Beyenbach, K.W., Skaer, H., and Dow, J.A.T. (2010). The developmental, molecular, and transport biology of Malpighian tubules. Annu Rev Entomol 55, 351–374.

Bilder, D., Li, M., and Perrimon, N. (2000). Cooperative regulation of cell polarity and growth by Drosophila tumor suppressors. Science 289, 113–116.

Bilder, D., Schober, M., and Perrimon, N. (2003). Integrated activity of PDZ protein complexes regulates epithelial polarity. Nat Cell Biol 5, 53–58.

Biteau, B., Hochmuth, C.E., and Jasper, H. (2008). JNK activity in somatic stem cells causes loss of tissue homeostasis in the aging Drosophila gut. Cell Stem Cell 3, 442–455.

Bonello, T.T., Choi, W., and Peifer, M. (2019). Scribble and Discs-large direct initial assembly and positioning of adherens junctions during the establishment of apical-basal polarity. Development 146, dev180976.

Cabrero, P., Terhzaz, S., Dornan, A.J., Ghimire, S., Holmes, H.L., Turin, D.R., Romero, M.F., Davies, S.A., and Dow, J.A.T. (2020). Specialized stellate cells offer a privileged route for rapid water flux in Drosophila renal tubule. Proc Natl Acad Sci USA 197, 201915943.

Cabrero, P., Terhzaz, S., Romero, M.F., Davies, S.A., Blumenthal, E.M., and Dow, J.A.T. (2014). Chloride channels in stellate cells are essential for uniquely high secretion rates in neuropeptide-stimulated Drosophila diuresis. Proc Natl Acad Sci USA 111, 14301–14306.

Clark, R.I., and Walker, D.W. (2018). Role of gut microbiota in aging-related health decline: insights from invertebrate models. Cell Mol Life Sci 75, 93–101.

Clark, R.I., Salazar, A., Yamada, R., Fitz-Gibbon, S., Morselli, M., Alcaraz, J., Rana, A., Rera, M., Pellegrini, M., Ja, W.W., et al. (2015). Distinct Shifts in Microbiota Composition during Drosophila Aging Impair Intestinal Function and Drive Mortality. Cell Rep 12, 1656–1667.

Cohen, E., Sawyer, J.K., Peterson, N.G., Dow, J.A.T., and Fox, D.T. (2020). Physiology, Development, and Disease Modeling in the Drosophila Excretory System. Genetics 214, 235–264.

Davies, S.A., Overend, G., Sebastian, S., Cundall, M., Cabrero, P., Dow, J.A.T., and Terhzaz, S. (2012). Immune and stress response “cross-talk” in the Drosophila Malpighian tubule. J Insect Physiol 58, 488–497.

Denholm, B. (2013). Shaping up for action: the path to physiological maturation in the renal tubules of Drosophila. Organogenesis 9, 40–54.

Denholm, B., Hu, N., Fauquier, T., Caubit, X., Fasano, L., and Skaer, H. (2013). The tiptop/teashirt genes regulate cell differentiation and renal physiology in Drosophila. Development 140, 1100–1110.

Denholm, B., Sudarsan, V., Pasalodos-Sanchez, S., Artero, R., Lawrence, P.A., Maddrell, S., Baylies, M., and Skaer, H. (2003). Dual origin of the renal tubules in Drosophila: mesodermal cells integrate and polarize to establish secretory function. Curr Biol 13, 1052–1057.

Dow, J.A.T., K.A. Halberg, S. Terhzaz, and S.A. Davies. 2018. *Drosophila* as a Model for Neuroendocrine Control of Renal Homeostasis. *In* Model Animals in Neuroendocrinology. 81–100.

Dow, J.A.T., (2012). The versatile stellate cell –More than just a space-filler. J Insect Physiol 58, 467–472.

Dow, J.A.T., Maddrell, S.H., Görtz, A., Skaer, N.J., Brogan, S., and Kaiser, K. (1994). The malpighian tubules of Drosophila melanogaster: a novel phenotype for studies of fluid secretion and its control. J Exp Biol 197, 421–428.

Elsum, I., Yates, L., Humbert, P.O., and Richardson, H.E. (2012). The Scribble-Dlg-Lgl polarity module in development and cancer: from flies to man. Essays Biochem. 53, 141–168.

Genova, J.L., and Fehon, R.G. (2003). Neuroglian, Gliotactin, and the Na+/K+ ATPase are essential for septate junction function in Drosophila. J. Cell Biol. 161, 979–989.

Halberg, K.A., S.M. Rainey, I.R. Veland, H. Neuert, A.J. Dornan, C. Klambt, S.-A. Davies, and J.A.T. Dow. 2016. The cell adhesion molecule Fasciclin2 regulates brush border length and organization in Drosophila renal tubules. Nat Commun. 7.

Halberg, K.A., Terhzaz, S., Cabrero, P., Davies, S.A., and Dow, J.A.T., (2015). Tracing the evolutionary origins of insect renal function. Nat Comms 6, 6800.

Harris, T.J.C., and Peifer, M. (2004). Adherens junction-dependent and - independent steps in the establishment of epithelial cell polarity in Drosophila. J. Cell Biol. 167, 135–147.

Hu, D.-J.K., and Jasper, H. (2017). Epithelia: Understanding the Cell Biology of Intestinal Barrier Dysfunction. Curr Biol 27, R185–R187.

Humbert, P.O., Grzeschik, N.A., Brumby, A.M., Galea, R., Elsum, I., and Richardson, H.E. (2008). Control of tumourigenesis by the Scribble/Dlg/Lgl polarity module. Oncogene 27, 6888–6907.

Humbert, P., Russell, S., and Richardson, H. (2003). Dlg, Scribble and Lgl in cell polarity, cell proliferation and cancer. Bioessays 25, 542–553.

Izumi, Y., and Furuse, M. (2014). Molecular organization and function of invertebrate occluding junctions. Semin Cell Dev Biol.

Izumi, Y., Furuse, K., and Furuse, M. (2019). Septate junctions regulate gut homeostasis through regulation of stem cell proliferation and enterocyte behavior in Drosophila. J Cell Sci jcs.232108.

Izumi, Y., Motoishi, M., Furuse, K., and Furuse, M. (2016). A tetraspanin regulates septate junction formation in Drosophila midgut. J Cell Sci 129, 1155–1164.

Izumi, Y., Yanagihashi, Y., and Furuse, M. (2012). A novel protein complex, Mesh-Ssk, is required for septate junction formation in the Drosophila midgut. J Cell Sci 125, 4923–4933.

Jonusaite, S., Beyenbach, K.W., Meyer, H., Paululat, A., Izumi, Y., Furuse, M., and Rodan, A.R. (2020). The septate junction protein Mesh is required for epithelial morphogenesis, ion transport, and paracellular permeability in the Drosophila Malpighian tubule. Am J Physiol, Cell Physiol 318, C675–C694.

Khoury, M.J., and Bilder, D. (2020). Distinct activities of Scrib module proteins organize epithelial polarity. Proc Natl Acad Sci USA 430, 201918462.

Lamb, R.S., Ward, R.E., Schweizer, L., and Fehon, R.G. (1998). Drosophila coracle, a member of the protein 4.1 superfamily, has essential structural functions in the septate junctions and developmental functions in embryonic and adult epithelial cells. Mol. Biol. Cell 9, 3505–3519.

Lane, N.J., and Skaer, H.I. (2016). Intercellular Junctions in Insect Tissues. Sciencedirect.com.

Martínez-Corrales, G., Cabrero, P., Dow, J.A.T., Terhzaz, S., and Davies, S.A. (2019). Novel roles for GATAe in growth, maintenance and proliferation of cell populations in the Drosophila renal tubule. Development dev.178087.

McGee, M.D., Weber, D., Day, N., Vitelli, C., Crippen, D., Herndon, L.A., Hall, D.H., and Melov, S. (2011). Loss of intestinal nuclei and intestinal integrity in aging C. elegans. Aging Cell 10, 699–710.

McGuire, S.E., Mao, Z., and Davis, R.L. (2004). Spatiotemporal gene expression targeting with the TARGET and gene-switch systems in Drosophila. Sci. STKE 2004, pl6.

Nelson, W.J. (2003). Adaptation of core mechanisms to generate cell polarity. Nature 422, 766–774.

Noirot-Timothee, C., and Noirot, C. (1980). Septate and scalariform junctions in arthropods. Int. Rev. Cytol. 63, 97–140.

Oshima, K., and Fehon, R.G. (2011). Analysis of protein dynamics within the septate junction reveals a highly stable core protein complex that does not include the basolateral polarity protein Discs large. J Cell Sci 124, 2861–2871.

Patrick, M.L., Aimanova, K., Sanders, H.R., and Gill, S.S. (2006). P-type Na+/K+-ATPase and V-type H+-ATPase expression patterns in the osmoregulatory organs of larval and adult mosquito Aedes aegypti. J Exp Biol 209, 4638–4651.

Regan, J.C., Khericha, M., Dobson, A.J., Bolukbasi, E., Rattanavirotkul, N., and Partridge, L. (2016). Sex difference in pathology of the ageing gut mediates the greater response of female lifespan to dietary restriction. Elife 5, e10956.

Rera, M., Azizi, M.J., and Walker, D.W. (2013a). Organ-specific mediation of lifespan extension: more than a gut feeling? Ageing Res. Rev. 12, 436–444.

Rera, M., Clark, R.I., and Walker, D.W. (2012). Intestinal barrier dysfunction links metabolic and inflammatory markers of aging to death in Drosophila. Proc Natl Acad Sci USA 109, 21528–21533.

Rera, M., Clark, R.I., and Walker, D.W. (2013b). Why do old flies die? Aging (Albany NY) 5, 586–587.

Resnik-Docampo, M., Koehler, C.L., Clark, R.I., Schinaman, J.M., Sauer, V., Wong, D.M., Lewis, S., D’Alterio, C., Walker, D.W., and Jones, D.L. (2017). Tricellular junctions regulate intestinal stem cell behaviour to maintain homeostasis. Nat Cell Biol 19, 52–59.

Resnik-Docampo, M., Sauer, V., Schinaman, J.M., Clark, R.I., Walker, D.W., and Jones, D.L. (2018). Keeping it tight: The relationship between bacterial dysbiosis, septate junctions, and the intestinal barrier in Drosophila. Fly (Austin) 12, 34–40.

Rose, M.R., Flatt, T., Graves, J.L., Greer, L.F., Martinez, D.E., Matos, M., Mueller, L.D., Shmookler Reis, R.J., and Shahrestani, P. (2012). What is Aging? Front Genet 3, 134.

Royer, C., and Lu, X. (2011). Epithelial cell polarity: a major gatekeeper against cancer? Cell Death Differ. 18, 1470–1477.

Salazar, A.M., Resnik-Docampo, M., Ulgherait, M., Clark, R.I., Shirasu-Hiza, M., Jones, D.L., and Walker, D.W. (2018). Intestinal Snakeskin Limits Microbial Dysbiosis during Aging and Promotes Longevity. iScience 9, 229–243.

Singh, S.R., and Hou, S.X. (2008). Lessons learned about adult kidney stem cells from the malpighian tubules of Drosophila. J Am Soc Nephrol 19, 660–666.

Singh, S.R., and Hou, S.X. (2009). Multipotent stem cells in the Malpighian tubules of adult Drosophila melanogaster. J Exp Biol 212, 413–423.

Singh, S.R., Zeng, X., Zheng, Z., and Hou, S.X. (2011). The adult Drosophila gastric and stomach organs are maintained by a multipotent stem cell pool at the foregut/midgut junction in the cardia (proventriculus). Cell Cycle 10, 1109–1120.

Skaer, H. 1993. The alimentary canal. *In* The Development of Drosophila melanogaster. Vol. 2. M. Bate and A. Martinez Arias, editors. Cold Spring Harbor Press, Cold Spring Harbor. 941–1012.

Sözen, M.A., Armstrong, J.D., Yang, M., Kaiser, K., and Dow, J.A.T., (1997). Functional domains are specified to single-cell resolution in a Drosophila epithelium. Proc Natl Acad Sci USA 94, 5207–5212.

Su, W.-H., Mruk, D.D., Wong, E.W.P., Lui, W.-Y., and Cheng, C.Y. (2012). Polarity protein complex Scribble/Lgl/Dlg and epithelial cell barriers. Adv. Exp. Med. Biol. 763, 149–170.

Takashima, S., Paul, M., Aghajanian, P., Younossi-Hartenstein, A., and Hartenstein, V. (2013). Migration of Drosophila intestinal stem cells across organ boundaries. Development 140, 1903–1911.

Tepass, U., and Hartenstein, V. (1994). The development of cellular junctions in the Drosophila embryo. Dev Biol 161, 563–596.

Tepass, U., Tanentzapf, G., Ward, R., and Fehon, R. (2001). Epithelial cell polarity and cell junctions in Drosophila. Annu Rev Genet 35, 747–784.

Terhzaz, S., Finlayson, A.J., Stirrat, L., Yang, J., Tricoire, H., Woods, D.J., Dow, J.A.T., and Davies, S.A. (2010). Cell-specific inositol 1,4,5 trisphosphate 3-kinase mediates epithelial cell apoptosis in response to oxidative stress in Drosophila. Cell Signal 22, 737–748.

Terhzaz, S., O’Connell, F.C., Pollock, V.P., Kean, L., Davies, S.A., Veenstra, J.A., and Dow, J.A.T., (1999). Isolation and characterization of a leucokinin-like peptide of Drosophila melanogaster. J Exp Biol 202, 3667–3676.

Texada, M.J., A.F. Jorgensen, C.F. Christensen, T. Koyama, A. Malita, D.K. Smith, D.F.M. Marple, E.T. Danielsen, S.K. Petersen, J.L. Hansen, K.A. Halberg, and K.F. Rewitz. 2019. A fat-tissue sensor couples growth to oxygen availability by remotely controlling insulin secretion. Nat Commun. 10:1955.

Verma, P., and M.G. Tapadia. 2014. Epithelial immune response in Drosophila malpighian tubules: interplay between Diap2 and ion channels. Journal of cellular physiology. 229:1078–1095.

Wang, C., and Spradling, A.C. (2020). An abundant quiescent stem cell population in Drosophila Malpighian tubules protects principal cells from kidney stones. Elife 9, 19.

Wodarz, A., and Näthke, I. (2007). Cell polarity in development and cancer. Nat Cell Biol 9, 1016–1024.

Woods, D.F., Hough, C., Peel, D., Callaini, G., and Bryant, P.J. (1996). Dlg protein is required for junction structure, cell polarity, and proliferation control in Drosophila epithelia. J. Cell Biol. 134, 1469–1482.

Wu, V.M., Schulte, J., Hirschi, A., Tepass, U., and Beitel, G.J. (2004). Sinuous is a Drosophila claudin required for septate junction organization and epithelial tube size control. J. Cell Biol. 164, 313–323.

Xu, C., Tang, H.-W., Hung, R.-J., Hu, Y., Ni, X., Housden, B.E., and Perrimon, N. (2019). The Septate Junction Protein Tsp2A Restricts Intestinal Stem Cell Activity via Endocytic Regulation of aPKC and Hippo Signaling. Cell Rep 26, 670–688.e676.

Yanagihashi, Y., Usui, T., Izumi, Y., Yonemura, S., Sumida, M., Tsukita, S., Uemura, T., and Furuse, M. (2012). Snakeskin, a membrane protein associated with smooth septate junctions, is required for intestinal barrier function in Drosophila. J Cell Sci 125, 1980–1990.

Yang, J., McCart, C., Woods, D.J., Terhzaz, S., Greenwood, K.G., ffrench-Constant, R.H., and Dow, J.A.T., (2007). A Drosophila systems approach to xenobiotic metabolism. Physiol Genomics 30, 223–231.

Yu, J.C., and Fernandez-Gonzalez, R. (2016). Local mechanical forces promote polarized junctional assembly and axis elongation in Drosophila. Elife 5.

