## Supplemental Figures S1-S8 for "The septate junction protein Snakeskin is critical for epithelial barrier function and tissue homeostasis in the Malpighian tubules of adult *Drosophila*"

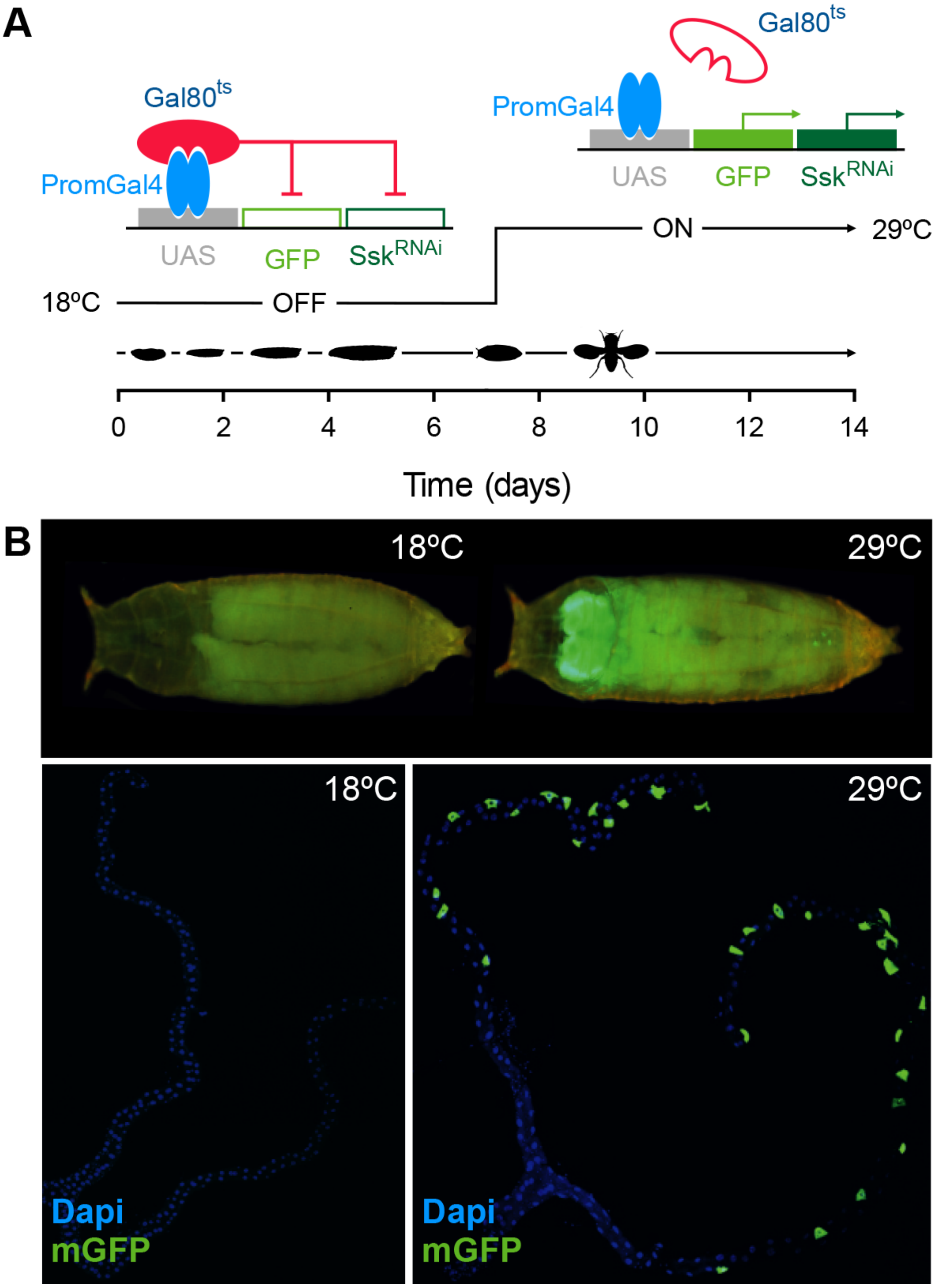


**Figure S1. Stratagem for cell-specific spatio-temporal knockdown of *Ssk* expression in adult Malpighian tubules**

(A) Temperature-sensitive stratagem for gene silencing of *Snakeskin* specifically in the stellate (SC) or principal (PC) cells of adult Malpighian tubules

(MTs) using the *c724*GAL4 and *Uro*GAL4 drivers respectively. GAL4 (PromGAL4) driver expression is repressed at the permissive temperature (18ºC) by expression of tubPGAL80$^{ts}$, precluding expression of GAL4-responsive UAS-transgenes (*UAS-mCD8::GFP* and *UAS-Ssk*$^{RNAi}$; red line). Experimental animals are transferred at late L3/white pre-pupal stage to the restrictive temperature (29ºC), repressing tubPGAL80$^{ts}$ expression, allowing expression of the GAL4-responsive UAS-transgenes in a spatially restricted manner (green arrows). (B) Membrane-bound green fluorescent protein (mGFP) expression absent in control (raised at the permissive temperature, 18ºC) but evident in experimental (raised at the restrictive temperature, 29ºC) 24 hr pupae and stellate cells of 5 D adult Malpighian tubules in SC$^{Ssk\mathrm{RNAi}}$ flies.

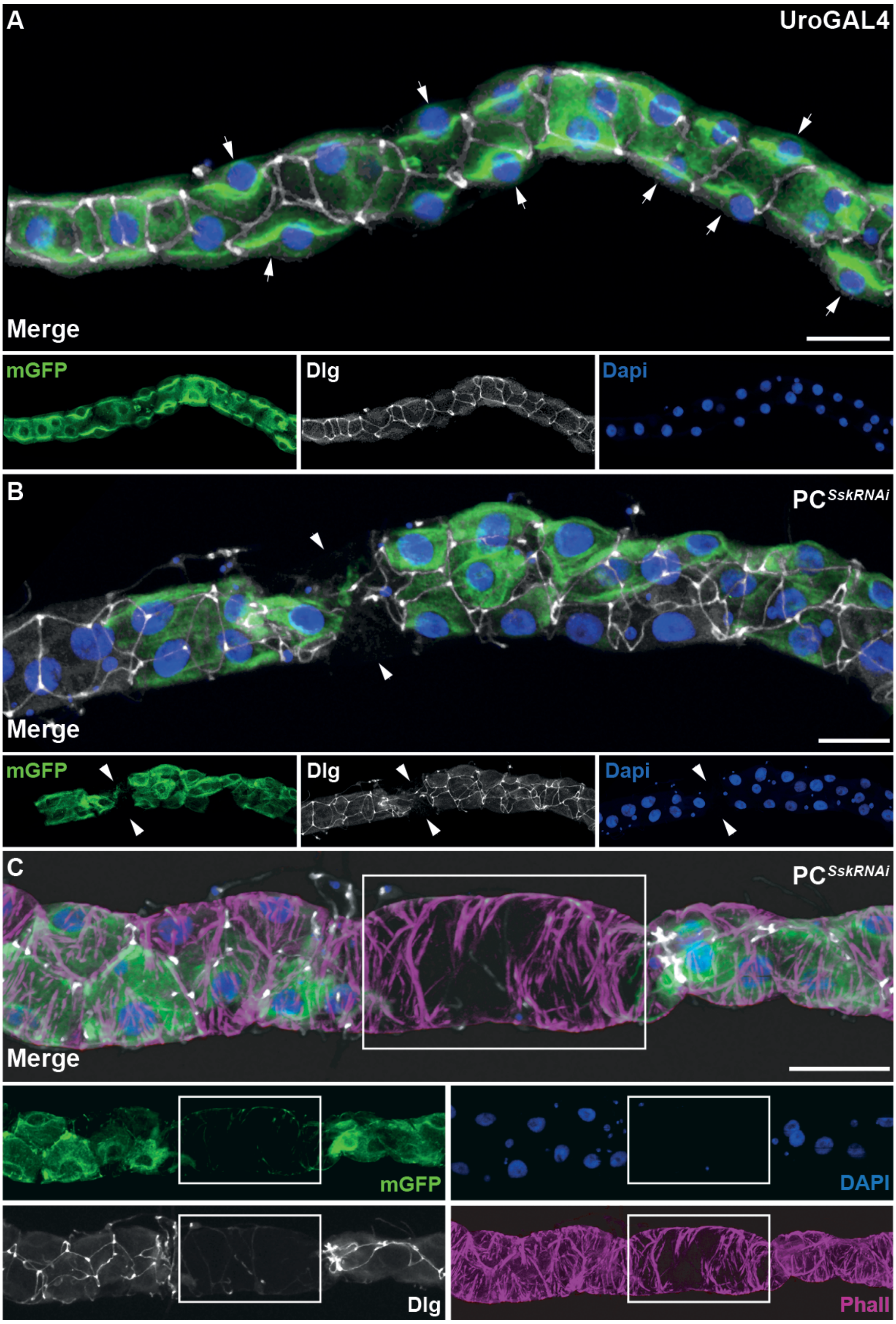


**Figure S2. Cell-specific depletion of *Ssk* in adult Malpighian tubules results in loss of cellular, junctional and cytoarchitectural organisation.** (A) *Uro*GAL4 driving membrane-bound green fluorescent protein (mGFP) in 5 day adult MT. Expression in PCs exhibiting stereotypical stellar morphology, with smoothly organised junctions throughout realized by anti-discs large (Dlg). mGFP expression exhibiting apical bias of expression (arrows). (B) 5 day experimental adult PC$^{Ssk\text{RNAi}}$ MT. PCs exhibit disorganised, or missing, junctional complexes and apparent loss of apical bias in mGFP expression. Complete absence of staining indicating apoptotic cells indicated (arrowheads). (C) 5 day experimental adult PC$^{Ssk\text{RNAi}}$ MT exhibiting absence of cellular architecture, realised by Phalloidin (Phall) staining as well as junctional expression, mGFP and Dapi staining (white box). mGFP, green; Phall, magenta; Dlg, white; Dapi, blue. Scale bars = 50 μm.

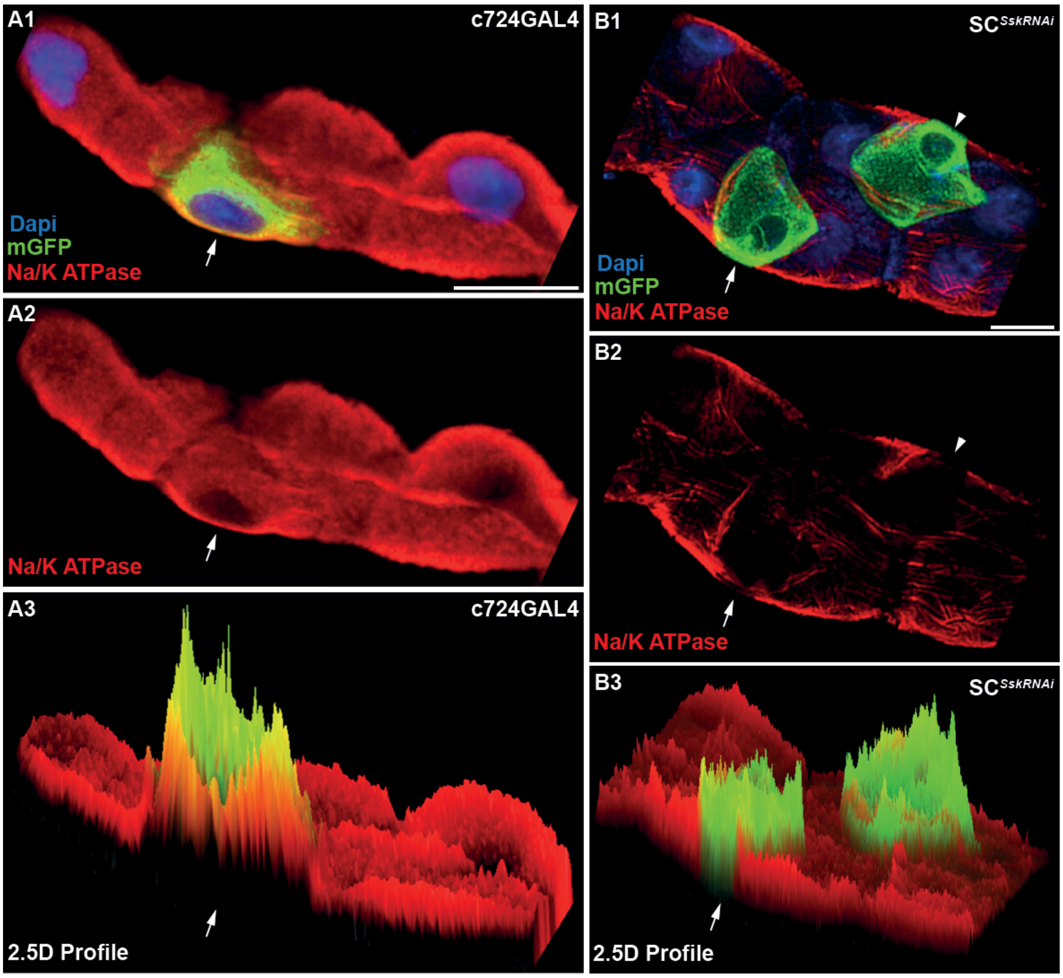


**Figure S3. cell-specific depletion of *Ssk* in adult Malpighian tubules results in loss of apicobasal polarity marker.**

(A) Subset of z-stack of *c724*GAL4 driving membrane-bound green fluorescent protein (mGFP) in 5 day adult MT. A1, merged image of confocal stack subset, exhibiting the basal staining for Na+/K+ alpha subunit (Na/K ATPase; arrow). A2, Na/K ATPase alone, highlighting expression at basal cell boundaries. A3, 2.5D rendering of A1 to illustrate Na/K ATPase expression biased to the basal cell boundary of SC (arrow). (B) Subset of z-stack of 5 day adult $SC^{\mathit{Ssk}\mathrm{RNAi}}$ MT. B1, merged image of confocal stack subset, detailing basal edge of SC exhibiting absence of staining for Na/K ATPase at the basal cell boundary (arrow). Second SC with absent Na/K ATPase at the basal cell boundary (arrowhead). B2, Na/K ATPase alone highlighting presence or absence of expression at basal cell boundaries (arrow and arrowhead). B3, 2.5D rendering B1 to illustrate absence of Na/K ATPase expression specific to mutant SC as compared with surrounding principal cells (arrow). Nb- both confocal stacks were collected using identical settings. mGFP, green; Na/K ATPase, red; Dapi, blue. Scale bars = 20 μm.

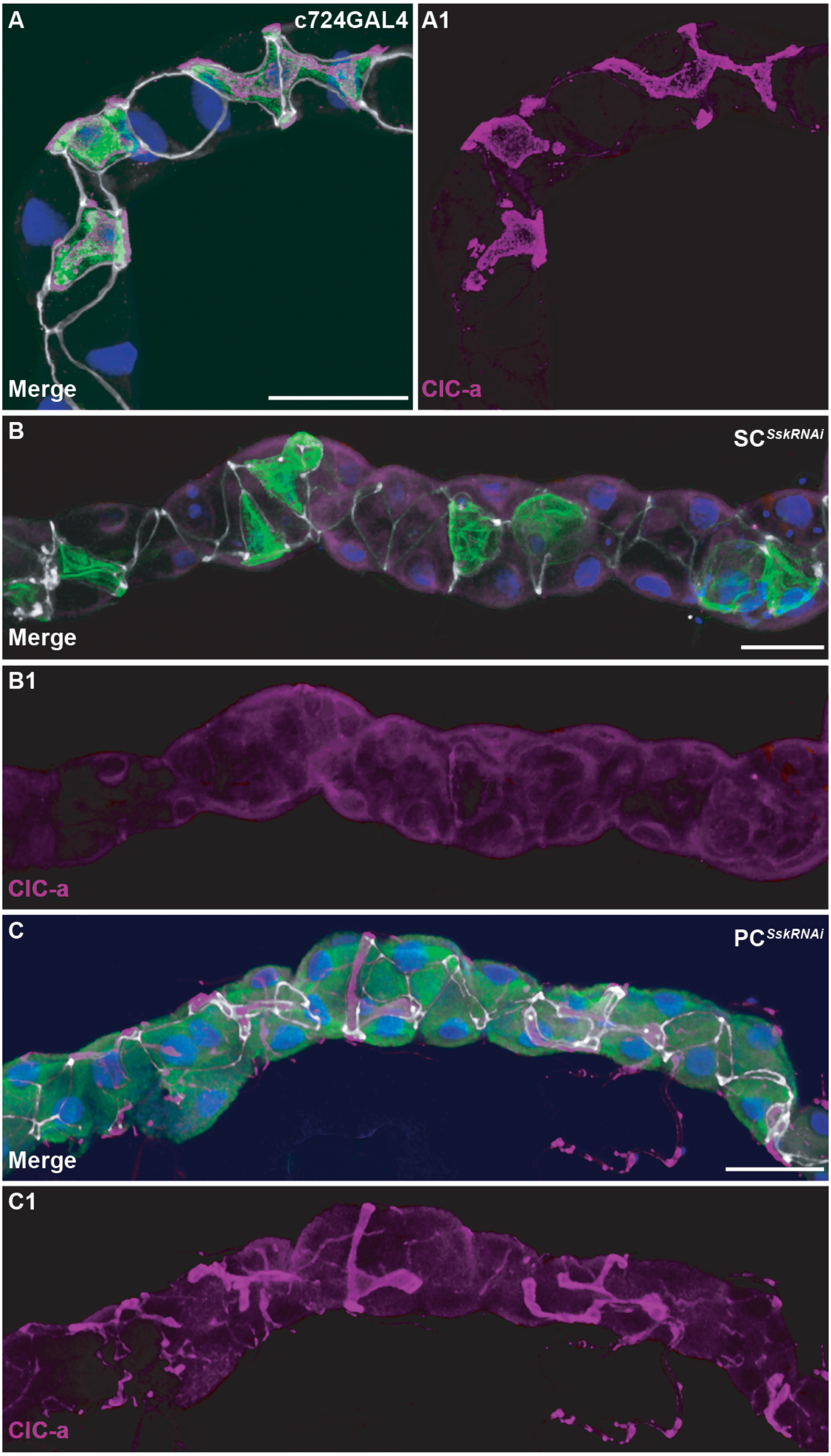
A
c724GAL4
Merge
A1
ClC-a
B
SC$^{SskRNAi}$
Merge
B1
ClC-a
C
PC$^{SskRNAi}$
Merge
C1
ClC-a

**Figure S4. SC-specific depletion of *Ssk* in adult Malpighian tubules results in loss of the SC-specific chloride channel-a.**

(A) *c724*GAL4 driving membrane-bound green fluorescent protein (mGFP) in 5 day adult MT, demonstrating co-expression with SC-specific anti-Chloride Channel-a **(**Clc-a). (B) 5 day adult SC$^{\mathit{Ssk}\mathrm{RNAi}}$ MT. SCs exhibit absence of SC-specific Clc-a expression. (C) 5 day adult PC$^{\mathit{Ssk}\mathrm{RNAi}}$ MT with with unaffected expression of Clc-a in SCs. Note SC stellar morphology is also unaffected. mGFP, green; Clc-A, magenta; Dlg, white; Dapi, blue. Scale bars = 50 $\mu$m.

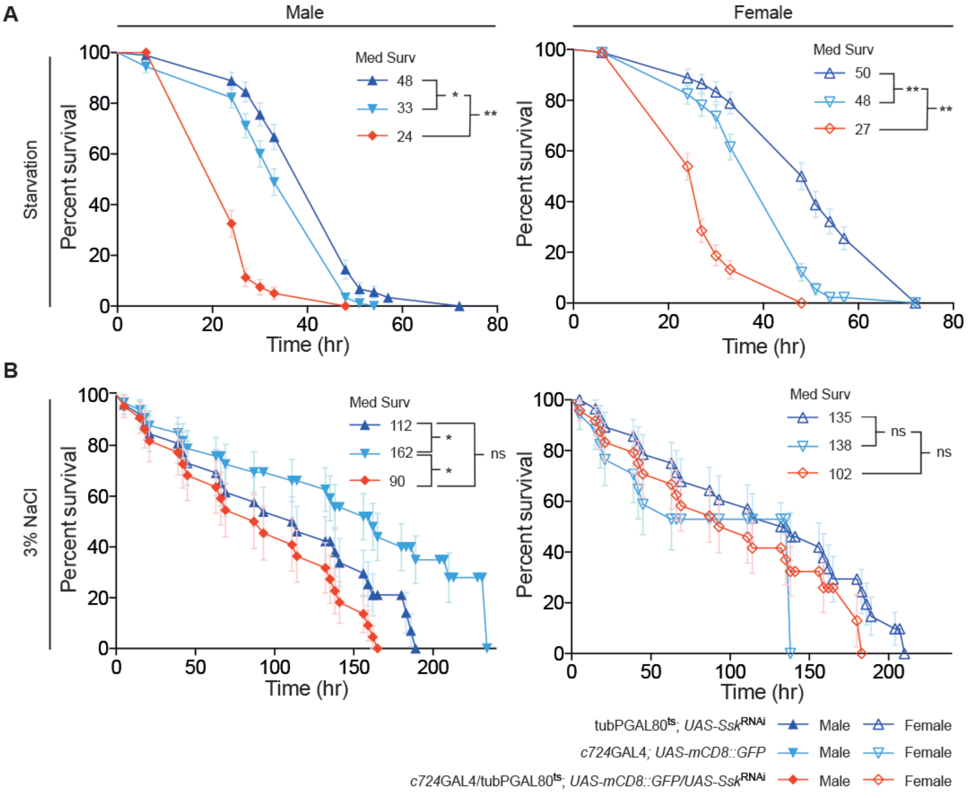


**Figure S5. SC-specific depletion of *Ssk* in adult Malpighian tubules results in sensitivity to starvation conditions but not salt stress.**

Both male and female 5 - 7 day old adult SC$^{\mathit{Ssk}\mathrm{RNAi}}$ flies demonstrate a significant reduction in viability during (A) (non-dessicating) starvation conditions compared to controls. This sensitivity is absent when these flies are challenged with (B) salt stress, with no significant difference in viability compared with controls. Mantel-Cox (Log rank) test. N's in parentheses. *P<0.05. **P<0.0001.

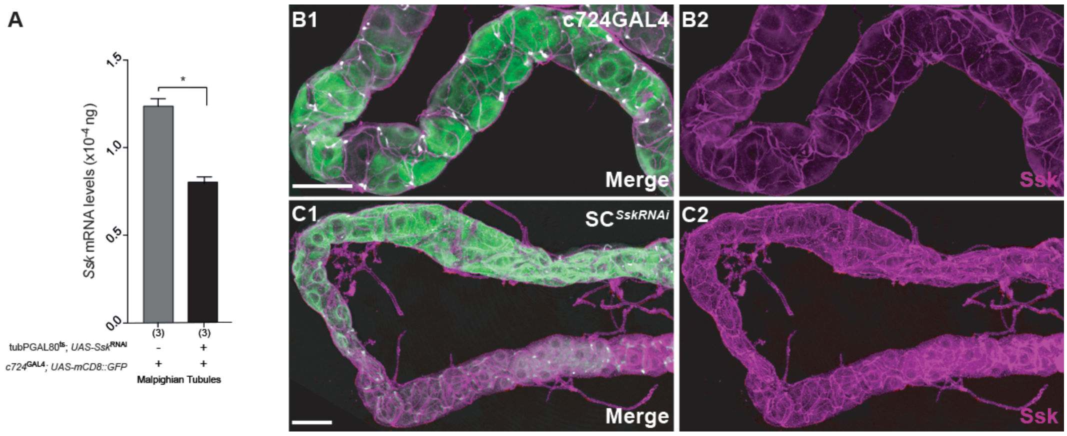


**Figure S6. Cell-specific depletion of *Ssk* results in hyperplasia of trachea supplying adult Malpighian tubules.**

(A) Quantitative RT-PCR analysis demonstrating ~35% knock down in *Ssk* mRNA expression levels in MTs between 5 day old adult SC$^{\mathit{Ssk}\mathrm{RNAi}}$ flies raised at the restrictive temperature (29ºC) compared with those raised at the permissive temperature (18ºC); 0.802 x $10^{-4}$ ng ± 0.038 vs 1.237 x $10^{-4}$ ng ± 0.043 mRNA respectively. (B) *Uro*GAL4 driving membrane-bound green fluorescent protein (mGFP) in the initial segment day of 5 D adult MT, demonstrating control levels of Ssk expression associated with junctions and, at lower levels, trachea. (C) Hyperplasia of trachea supplying MTs in adult PC$^{\mathit{Ssk}\mathrm{RNAi}}$ flies as realised by anti-Ssk. As with SC$^{\mathit{Ssk}\mathrm{RNAi}}$ flies, hyperplasia levels increase from initial to main segments. Nb- both confocal stacks were collected using identical settings. mGFP, green; Ssk, magenta; Dlg, white. Scale bars = 50 μm.

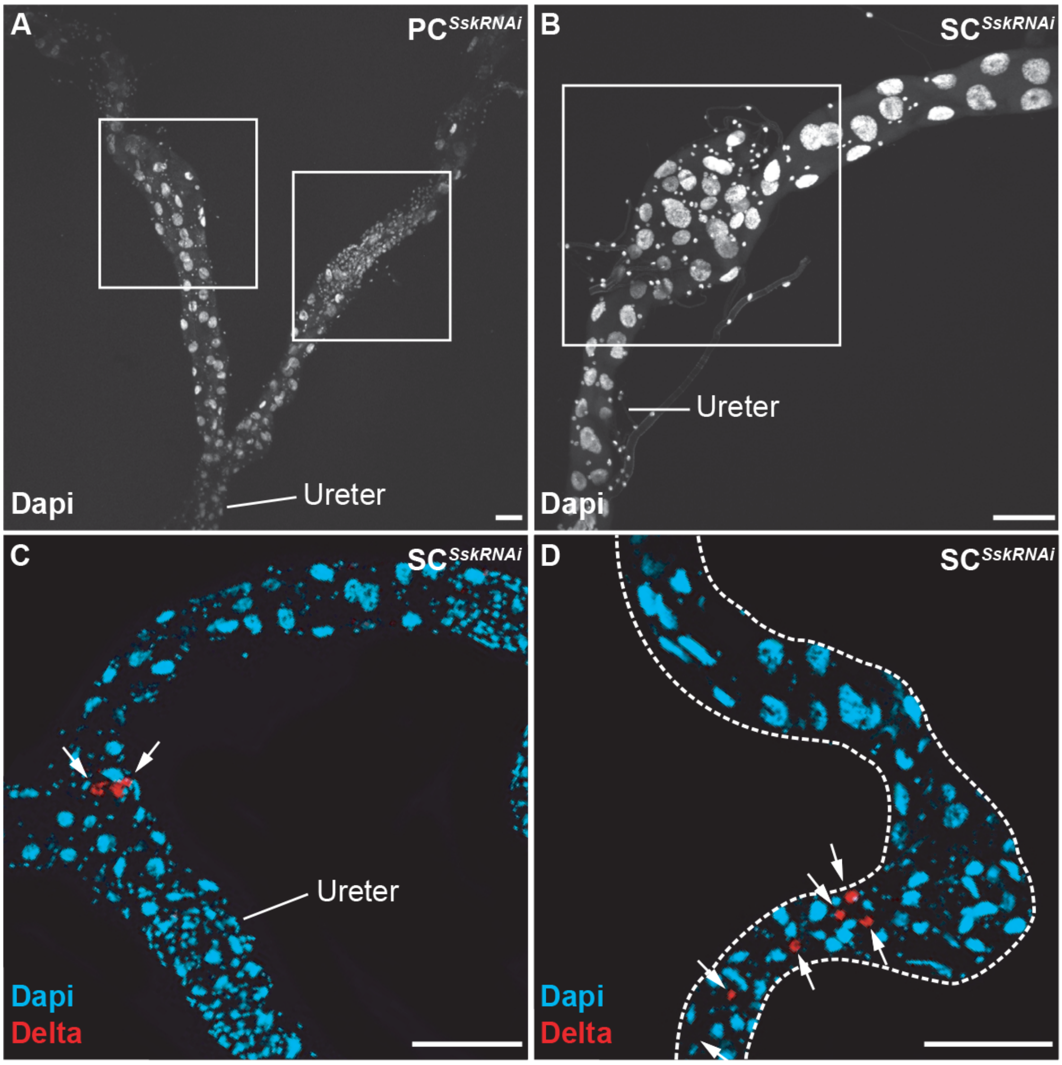


**Figure S7. Cell-specific depletion of *Ssk* results in cellular proliferation of tiny cells in adult Malpighian tubules.**

5 day PC$^{\mathit{Ssk}\mathrm{RNAi}}$ (A) and SC$^{\mathit{Ssk}\mathrm{RNAi}}$ flies (B) adult MTs exhibiting 'bulbar' areas with proliferation of 'tiny' (RNSC) cells nuclei (white boxes). Dapi, white. 5 day SC$^{\mathit{Ssk}\mathrm{RNAi}}$ adult MTs initial (C) and main (D) segments exhibiting co-expression of the proliferative cell marker anti-Delta (Delta) with 'tiny' (RNSC) cellsnuclei (arrows). Dapi, blue; Delta, red. Scale bars = 50 $\mu$m.

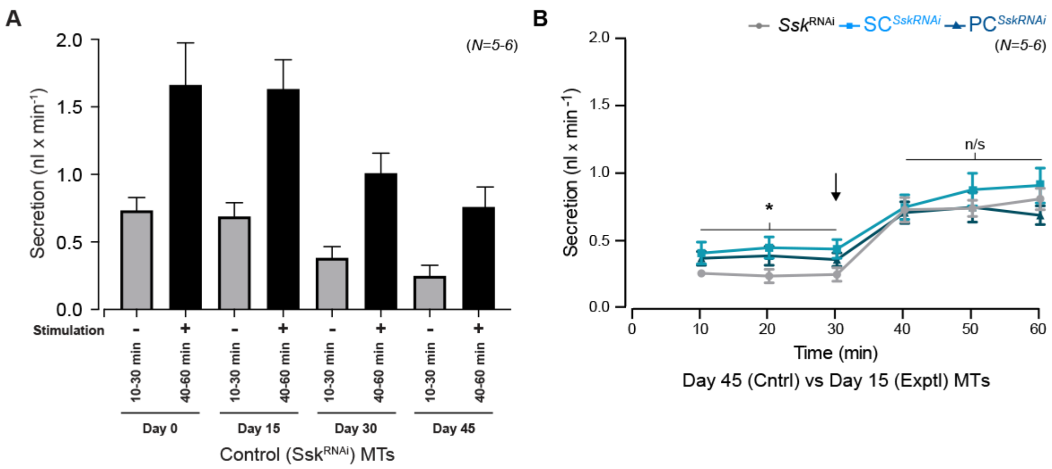


**Figure S8. Cell-specific *Ssk* depletion accelerates the age-related decline in function in adult Malpighian tubules.**

(A) Graph of basal (grey) and stimulated (black) secretion rates in control adult MTs over progressive time points, demonstrating the natural decline in tubule functional capacity that occurs as flies age. (B) Comparison of fluid secretion rates between 45 day old control and 15 day old $PC^{Ssk\text{RNAi}}$ and $SC^{Ssk\text{RNAi}}$ flies experimental adult MTs. While experimental MTs basal secretion rates are marginally better, there is no significant difference in secretion rates upon stimulation with Dromekinin (black arrow). *P<0.05, paired samples t-test. N's in parentheses.
